# CD4+CCR6+ T cells dominate the BCG-induced transcriptional signature

**DOI:** 10.1101/2021.08.20.457054

**Authors:** Akul Singhania, Paige Dubelko, Rebecca Kuan, William D. Chronister, Kaylin Muskat, Jyotirmoy Das, Elizabeth J. Phillips, Simon A. Mallal, Grégory Seumois, Pandurangan Vijayanand, Alessandro Sette, Maria Lerm, Bjoern Peters, Cecilia S. Lindestam Arlehamn

## Abstract

The century-old *Mycobacterium bovis* Bacillus Calmette-Guerin (BCG) remains the only licensed vaccine against tuberculosis (TB). Despite this, there is still a lot to learn about the immune response induced by BCG, both in terms of phenotype and specificity. Here, we investigated immune responses in adult individuals pre and 8 months post BCG vaccination. We specifically determined changes in gene expression, cell subset composition, DNA methylome, and the TCR repertoire induced in PBMCs and CD4 memory T cells associated with antigen stimulation by either BCG or a *Mycobacterium tuberculosis* (*Mtb*)-derived peptide pool. Following BCG vaccination, we observed increased frequencies of CCR6+ CD4 T cells, which includes both Th1* and Th17 subsets, and mucosal associated invariant T cells (MAITs). A large number of immune response genes and pathways were upregulated post BCG vaccination with similar patterns observed in both PBMCs and memory CD4 T cells, thus suggesting a substantial role for CD4 T cells in the cellular response to BCG. These upregulated genes and associated pathways were also reflected in the DNA methylome. We described both qualitative and quantitative changes in the BCG-specific TCR repertoire post vaccination, and importantly found evidence for similar TCR repertoires across different subjects. The immune signatures defined herein can be used to track and further characterize immune responses induced by BCG, and can serve as reference for benchmarking novel vaccination strategies.

## INTRODUCTION

Tuberculosis (TB), claims over 1.5 million lives every year, and is caused by infection with *Mycobacterium tuberculosis* (*Mtb*). The Bacillus Calmette-Guerin (BCG) vaccine, first introduced a century ago, remains the only approved vaccine against TB and most widely used vaccine in the world. BCG offers variable efficacy against pulmonary TB in all age-groups, but has high efficacy against severe forms of TB in young children ^1-3^. The underlying cause for this variable efficacy is difficult to pinpoint, and is likely due to multiple factors including geographical location and exposure to environmental mycobacteria. Many approaches have been considered to improve the BCG efficacy, including changes in the route of administration from the current intradermal standard ^4-7^, booster vaccinations with either BCG ^8-13^, or with *Mtb*-derived antigens ^14-16^, and recombinant BCG strains ^14,17^. These results have been mixed and a clear enhanced efficacy has not been achieved. Further rational vaccine improvement efforts are hindered by our incomplete understanding of the mechanisms of immune protection elicited by BCG.

BCG was developed empirically more than a century ago ^18^, and yet surprisingly little is known about BCG-induced immune responses to this date. The specific cell subset or subsets responsible for mediating BCG’s protective effects have not been clarified. While BCG-specific T cell reactivity does not mediate protection alone, it can be used as an immune correlate of *Mtb* infection and disease risk ^19,20^. An increased level of BCG-specific cells post-vaccination is frequently reported, but the BCG-specific T cell response has varied considerably between studies ^1,18,21,22^. Thus, studying the cellular response induced by BCG vaccination is a key component of understanding how it mediates protection and what immune responses are triggered.

Systems biology is a compelling approach that can be used to dissect the heterogeneity of immune responses following various perturbations (vaccination, infection, disease, autoimmunity, etc.). The resulting gene signatures and cellular profiles have proven of significant diagnostic, prognostic, or mechanistic value ^23^. Similar to what we have previously described for individuals with latent TB infection (LTBI) as compared to TB-uninfected controls [63, 64], here we used a comprehensive systems biology/multiomics approach to determine a global picture of the immune responses triggered by BCG vaccination in humans. We conducted a longitudinal study of immune responses associated with BCG vaccination in a cohort of BCG naïve adults. This allowed the opportunity to study cell subset changes, gene signatures and the TCR repertoire changes induced by BCG vaccination. The results indicate that BCG induced gene signatures in adults are primarily driven by the expansion of Th1* CD4 memory T cells and results in both qualitative and quantitative TCR repertoire changes.

## RESULTS

### The frequency of specific T cell subsets increases following BCG vaccination

To determine the effects of BCG vaccination on specific T cell responses and gene expression, PBMC samples were obtained from BCG naïve individuals (pre-vaccination). Subsequently, these individuals were administered the BCG vaccine, and PBMC samples were collected again 8 months after intradermal administration of BCG (post-vaccination), chosen as a representative of a time point were immune responses are expected to have reached a steady state memory phase (**Figure 1a**). PBMCs from both time points (pre- and post-vaccination) were assayed by flow cytometry and in parallel stimulated for 24 hours *in vitro* with the vaccine itself (BCG) or media (unstim) to identify BCG-induced immune responses based on Fluorospot and RNA-sequencing (**Figure 1a**).

**FIGURE 1.**
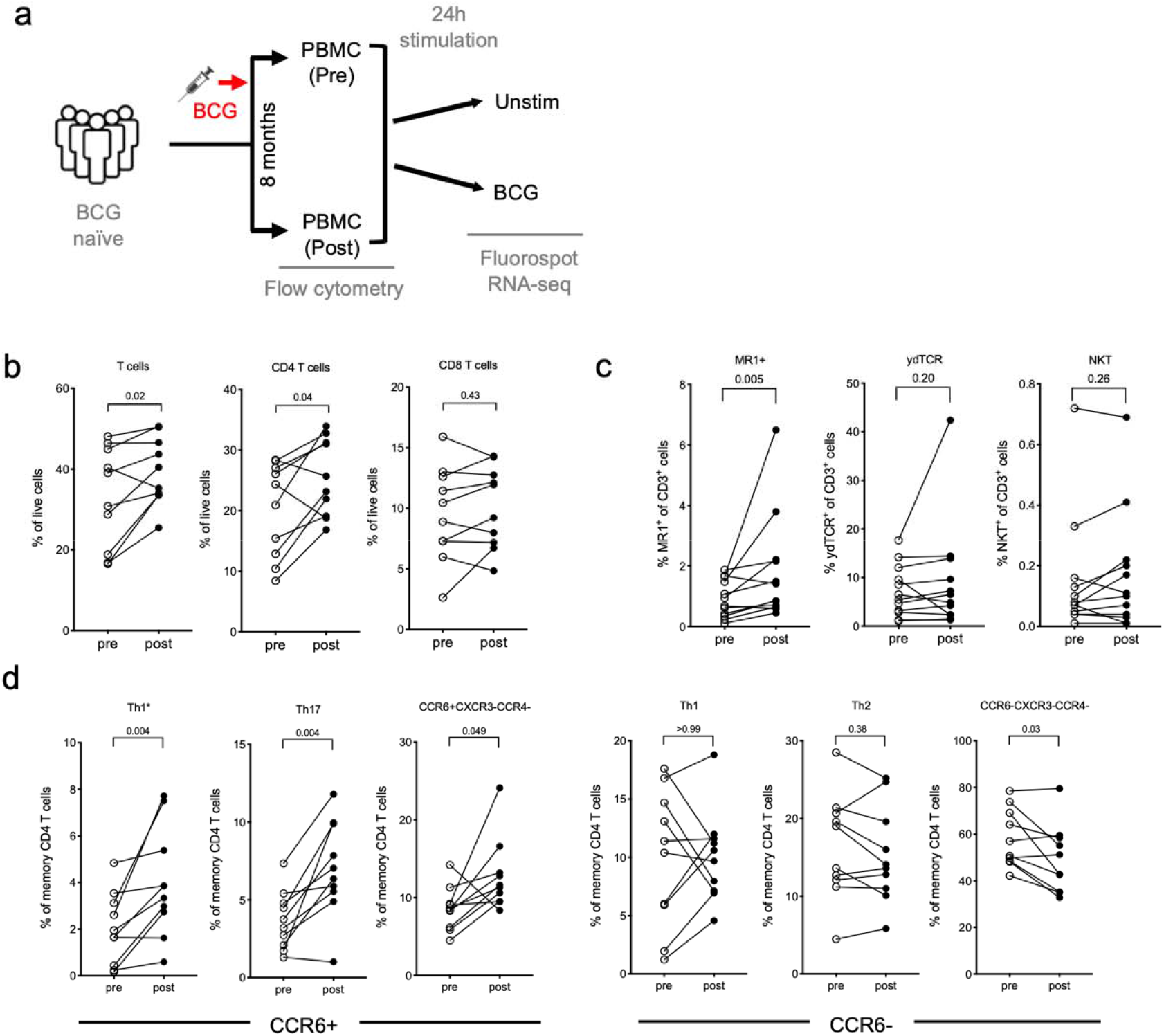
T cell subsets show an increase post BCG vaccination. **a** Schematic of the project workflow. Flow cytometry analysis was performed on PBMC samples obtained from BCG naïve individuals, pre and post BCG vaccination, and Fluorospot and RNA-sequencing on PBMCs stimulated with either DMSO or BCG for 24 hours. **b-d** Frequencies of cell subsets pre-(open circles) and post-(closed circles) BCG vaccination as determined by flow cytometry. Each point represents one participant, Wilcoxon matched pair signed rank test. **b** T cells, CD4 and CD8 T cells. **c** MAITs; as defined by staining with MR1 5-OP-RU tetramer, γδ T cells; defined by pan γδTCR, and NKTs; defined by Vα24 staining. **d** T-helper subsets defined by CXCR3, CCR6 and CCR4 expression.

We previously described a flow cytometry panel designed to quantitate the relative frequency of major PBMC subsets ^24^. In addition, we have previously identified increased frequencies of a specific Th subset, Th1* (CXCR3+CCR6+) ^25^, as well as a decrease in MR1+ T cells in individuals with latent TB infection (LTBI) as compared to TB negative subjects ^26^. Here, we investigated the frequency changes following BCG vaccination of major PBMC subsets and specific T cell subsets in the absence of antigen-specific stimulation. Upon BCG vaccination we observed an increase in T cell frequency, which was specifically driven by CD4 T cells while CD8 T cells remained unchanged (**Figure 1b**). In contrast, the frequency of B cells decreased following vaccination (**Supplementary figure 1a**). No changes were observed for monocytes, NK cells, or CD3+CD56+ T cell populations (**Supplementary figure 1a**). Memory CD4 and CD8 T cell populations, as defined by CCR7 and CD45RA expression, remained unchanged following vaccination (**Supplementary figure 1b, c**). In terms of non-conventional T cells, we found an increased frequency of MAITs (as defined by 5-OP-RU loaded MR1 tetramers) following vaccination, but no changes in γδ T cells and NKT cells (**Figure 1c**). Finally, we determined the frequencies of Th subsets, defined by CXCR3, CCR6 and CCR4 expression. All CCR6+ populations, including Th17 and Th1* cells, increased following BCG vaccination, whereas CCR6-populations, including Th1 and Th2 cells, remained unchanged (**Figure 1d**). These results are consistent with our previous identification of Th1* as the Th subset that contains the vast majority of *Mtb*-and non-tuberculous mycobacteria (NTM)-specific T cells ^25,27^, and the present results point towards a role of these subsets following BCG vaccination as well.

### BCG-induced T cell responses and gene expression changes are enhanced upon BCG vaccination

We next investigated the magnitude and quality of BCG-induced cellular immune responses. The magnitude of BCG-stimulated T cell responses, as measured by IFNγ Fluorospot assay, increased post-vaccination (**Figure 2a**). In comparison, no difference was observed in the magnitude of response against PHA, which was used as a positive control (**Figure 2a**).

**FIGURE 2.**
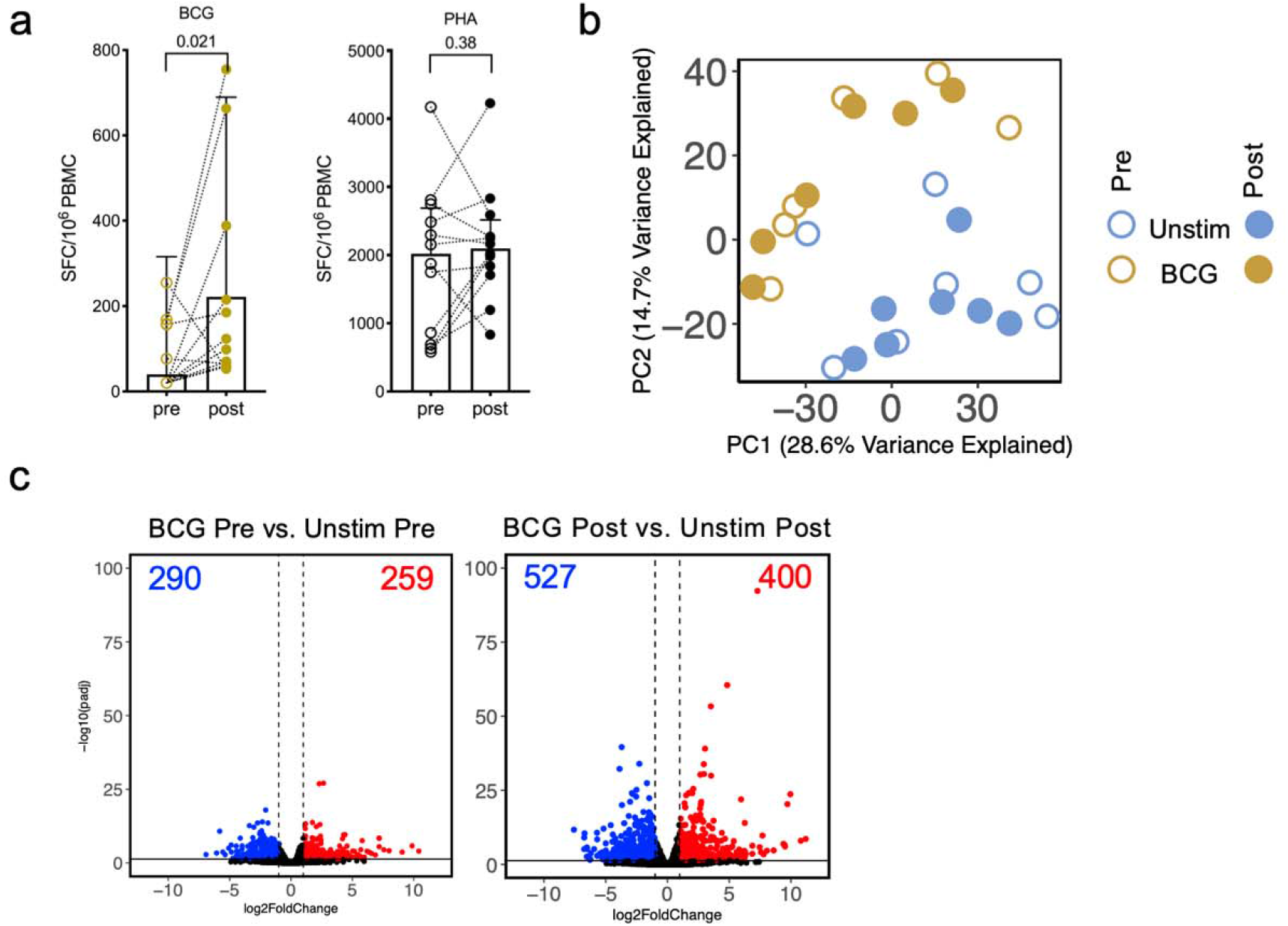
BCG-induced T cell responses and gene expression changes are enhanced upon BCG vaccination. **a** Paired point graphs depicting magnitude of IFNγ responses pre- and post-vaccination, against BCG and PHA as SFC per 10^6^ cultured PBMC as determined by Fluorospot. Each point and symbol represent one participant, median ± interquartile range is shown. Wilcoxon matched pair signed rank test. **b** PCA depicting the variation in global gene expression as a result of stimulation condition and vaccination. Pre-vaccination (open circles) and post vaccination (closed circles), unstimulated samples (blue) and BCG stimulated samples (gold). **c** Volcano plot depicting differentially expressed genes after BCG stimulation as compared to unstimulated samples, pre- and post-vaccination. Red indicates upregulated genes (adjusted p value < 0.05 and log_2_ fold change > 1) and blue indicates downregulated genes (adjusted p value < 0.05 and log_2_ fold change < -1).

To determine the effects of stimulation and vaccination on global gene expression, principal component analysis (PCA) was performed on the RNA-Seq data. The PCA showed a distinct separation of BCG stimulated and unstimulated samples (**Figure 2b**). In contrast, a smaller separation was observed in the pre- and post-vaccination samples, in each stimulated condition (**Figure 2b**).

In addition to the PCA analysis we performed differential gene expression analysis comparing BCG stimulated and unstimulated samples, to explore BCG-induced gene expression changes pre- and post-vaccination. Upon BCG vaccination, we observed an increase in the number of differentially expressed genes (DEG) (**Figure 2c; Supplementary table 1**). A total of 549 DEGs were identified when BCG-stimulated vs. unstimulated pre-vaccination samples were studied; 290 downregulated and 259 upregulated. This compares with a total of 927 DEGs identified post-vaccination; 527 downregulated and 400 upregulated. Importantly, BCG-induced T cell responses can be detected prior to vaccination, but they were enhanced 8 months post-vaccination.

### BCG-induced gene signatures are more pronounced after vaccination

We next investigated the BCG-induced gene signatures in more detail. We focused on the genes that were upregulated following BCG stimulation. A majority of the DEGs identified pre-vaccination were also differentially expressed post-vaccination (**Figure 3a**; 259 genes upregulated pre-vaccination and 400 genes upregulated post-vaccination, with an overlap of 175 genes). To determine if the unique DEGs identified both pre- and post-vaccination were truly unique, or if they were below the cut-off (padj values < 0.05 and log2 fold change > 1 or < - 1) for reaching significance in the other group, we compared fold changes for all upregulated genes identified pre- and post-vaccination. We observed that although most of these genes were upregulated both pre- and post-vaccination, the majority of the genes had higher fold changes post-vaccination (**Figure 3b**; **Supplementary table 1**; the 343 genes above the 45-degree slope represent those that were higher post-vaccination, and the 141 genes below the 45-degree slope were higher pre-vaccination). This suggests that although similar genes were upregulated upon BCG stimulation, the magnitude of this increase was greater post-vaccination.

**FIGURE 3.**
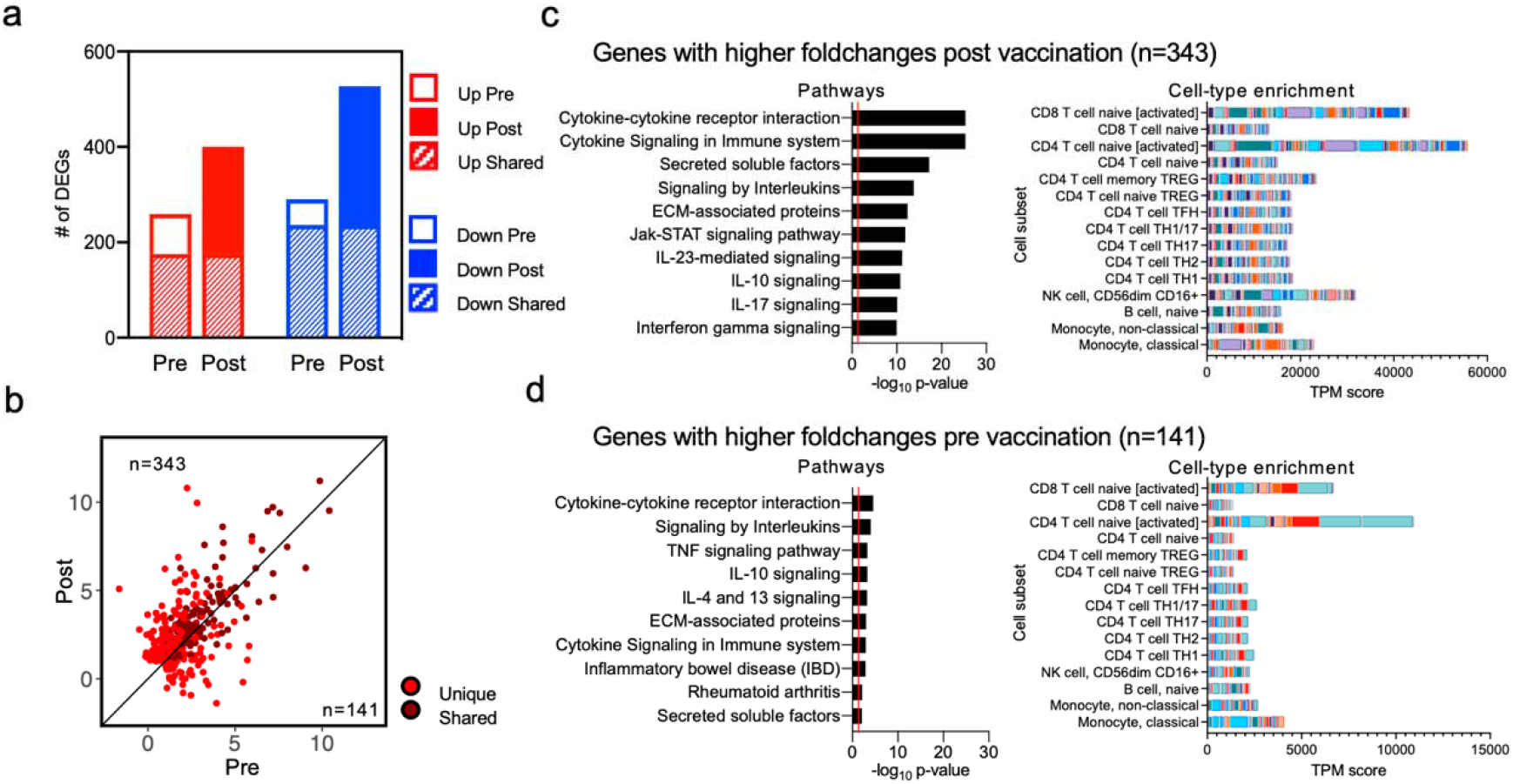
BCG-induced gene expression changes are enhanced upon BCG vaccination. **a** Bar plots representing the total number of differentially expressed genes pre- and post-vaccination. Empty and filled bars represent pre- and post-vaccination, respectively, color represents up or down regulation, and hatched filling represents overlap of genes between the pre and post time points. **b** Scatterplot of genes upregulated upon BCG stimulation in pre-vaccination (x-axis) and post-vaccination (y-axis). Each dot represents the log_2_ fold change for a gene, and color represents shared (gene significantly differentially expressed in both pre- and post-vaccination) or unique (differentially expressed in only one condition) expression. Black line at the 45° slope represents identical perturbation in pre- and post-vaccination, with genes above or below the line showing higher perturbation in the post or pre-group, respectively. **c, d** Pathway and cell-type enrichment (dice-database.org) for genes with **(c)** higher log_2_ fold changes post vaccination, and **(d)** higher log_2_ fold changes pre-vaccination. The ten most significant pathways are shown.

Several immune-related genes were upregulated in response to BCG stimulation both pre-and post-vaccination. These included chemokine ligands (CCL3, CCL4, CCL20, CXCL1, CXCL2, CXCL3, and CXCL9), T cell activation markers (CD38, CD69, and TNFRSF4 (OX40)), and cytokines (IFNG, IL1B, IL32, GZMB, and TNF), reflecting the broad immune response triggered by stimulation with BCG (**Supplementary table 1**). The genes that had a higher fold change post-vaccination included T cell activation genes (CD274 (PDL1) and DPP4 (CD26)), cytokines (IL13, IL17F, IL22, GZMA), chemokine ligands (CXCL5, CXCL8, CXCL13), and ligands for CXCR3 (CXCL10 (IP-10), and CXCL11), indicating a boosting of certain aspects of the BCG-induced immune response post-vaccination (**Supplementary table 1**).

Functional enrichment analysis of the upregulated genes showed that similar pathways, such as cytokine response, interleukin signaling and secreted factors, were identified in both the genes that were higher post-vaccination (**Figure 3c; Supplementary table 2**) and in genes higher pre-vaccination (**Figure 3d; Supplementary table 2**). However, as observed above, the magnitude of BCG related changes was greater post-vaccination and the common pathways identified in both groups had much greater significance post-vaccination. Moreover, pathways such as IL-23 mediated signaling, and IL-10 and IL-17 signaling were observed only in genes higher in post-vaccination (**Figure 3c**). Cell-type enrichment of upregulated genes showed an enrichment of activated CD4 and CD8 activated T cells in both groups (**Figure 3c and 3d**), and a slight enrichment of NK cells post-vaccination only (**Figure 3c**). These results demonstrate an increase in the magnitude of BCG-specific gene expression changes post BCG vaccination. Moreover, pathway enrichment suggested that a Th1*/Th17-like signature is associated with BCG vaccination, which was also observed by flow cytometry analysis.

### BCG-induced DNA methylation changes reflect the gene signatures

A higher magnitude of response upon microbial challenge is expected in epigenetically reprogrammed cells ^28^, and BCG has been shown to induce DNA methylation (DNAm) changes ^29^. Therefore, we next assessed the methylation status of >850,000 CpG sites in DNA derived from PBMC pre- and post-vaccination. First, using the Houseman algorithm to deconvolute cell types from PBMC using DNAm data ^30^, we found, similarly to the flow cytometry analysis an increased frequency of CD4 T cells post vaccination (**Supplementary figure 2a**). Unlike the PCA analysis for RNAseq of unstimulated PBMCs, the PCA analysis of the DNAm data revealed a clear separation between pre- and post-vaccination samples (**Figure 4a**). We next determined the differentially methylated CpG sites and identified 15,679 hypomethylated and 15,309 hypermethylated CpG sites (**Supplementary Figure 2b**), which were both primarily found in the gene bodies and intragenic regions (**Supplementary Figure 2c**). The CpG sites were mapped to genes to identify differentially methylated genes (DMGs), and a subsequent KEGG pathway analysis revealed the pathways affected by the DNAm changes (**Figure 4b**). We identified an IFNγ- and IL-17-related pathway (inflammatory bowel disease), a TNF- and CXCL10-related pathway (TNF signaling pathway), and a NOD2-like receptor signaling pathway. A comparison between the DMGs and the DEGs identified post-vaccination revealed that the majority of the hypo-methylated/upregulated genes were found in the identified pathways (**Figure 4c, d**). These results suggest a correlation between genes identified as upregulated in the RNAseq analysis and hypo-methylated genes.

**FIGURE 4.**
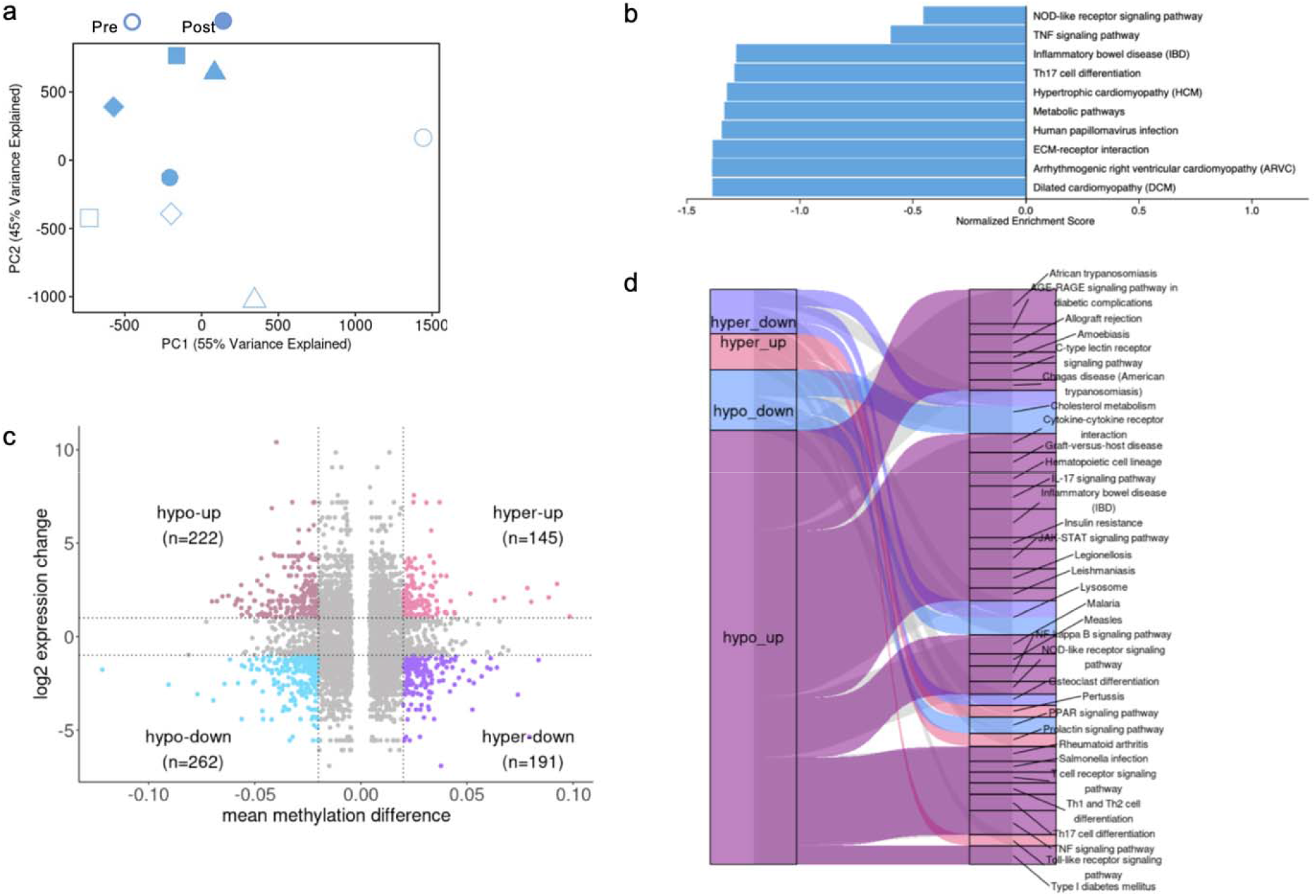
BCG-induced DNA methylation changes. **a** PCA of the overall DNA methylome data. Unstimulated pre-vaccination (open symbols) and post vaccination (closed symbols) samples. **b** Pathway enrichment for DMGs derived from hypomethylated CpGs. **c** Scatter plot representing the overlap of hypermethylated, hypomethylated, upregulated and downregulated genes post-vaccination. The vertical dotted lines represent the cut-off value of mmd ≥ |0.02| and the horizontal dotted lines represent the log2 expression value cut-off ≥ |1|. **d** Pathway enrichment analysis of overlapping significant DEGs and DMGs using the KEGG database The Gene Set Enrichment Analysis was performed using the 2000 permutations and BH-corrected p-value < 0.05 on the common gene list with mmd values. The resulting map of enriched pathways using KEGG database in different combinations of DMGs and DEGs is shown. Edge width indicates the number of significant pathways involved with the corresponding DMG-DEG list.

### MTB300-specific reactivity is not boosted by BCG vaccination

Next, we wanted to determine whether reactivity against a peptide pool defined in healthy *Mtb*-infected individuals, and with peptides homologous to peptides found in BCG, was boosted following BCG vaccination. PBMC samples from both time points (pre- and post-vaccination) were stimulated for 24h *in vitro* with the MTB300 peptide pool, as described above, and underwent Fluorospot analysis and RNA-Sequencing. The PCA revealed little separation between the MTB300 stimulated and unstimulated samples, suggesting stimulation with the MTB300 peptide pool does not have a large impact on overall global gene expression (**Figure 5a**). The lack of separation also suggested lower MTB300-specific reactivity as compared to that seen upon BCG stimulation. As previously noted, there was a smaller, but consistent, separation between pre- and post-vaccination samples within each stimulation condition **(Figure 5a)**. Moreover, while the magnitude of MTB300-specific IFNγ response increased post-vaccination (**Figure 5b**), it was lower than what was observed for BCG stimulation (**Figure 2a**).

**FIGURE 5.**
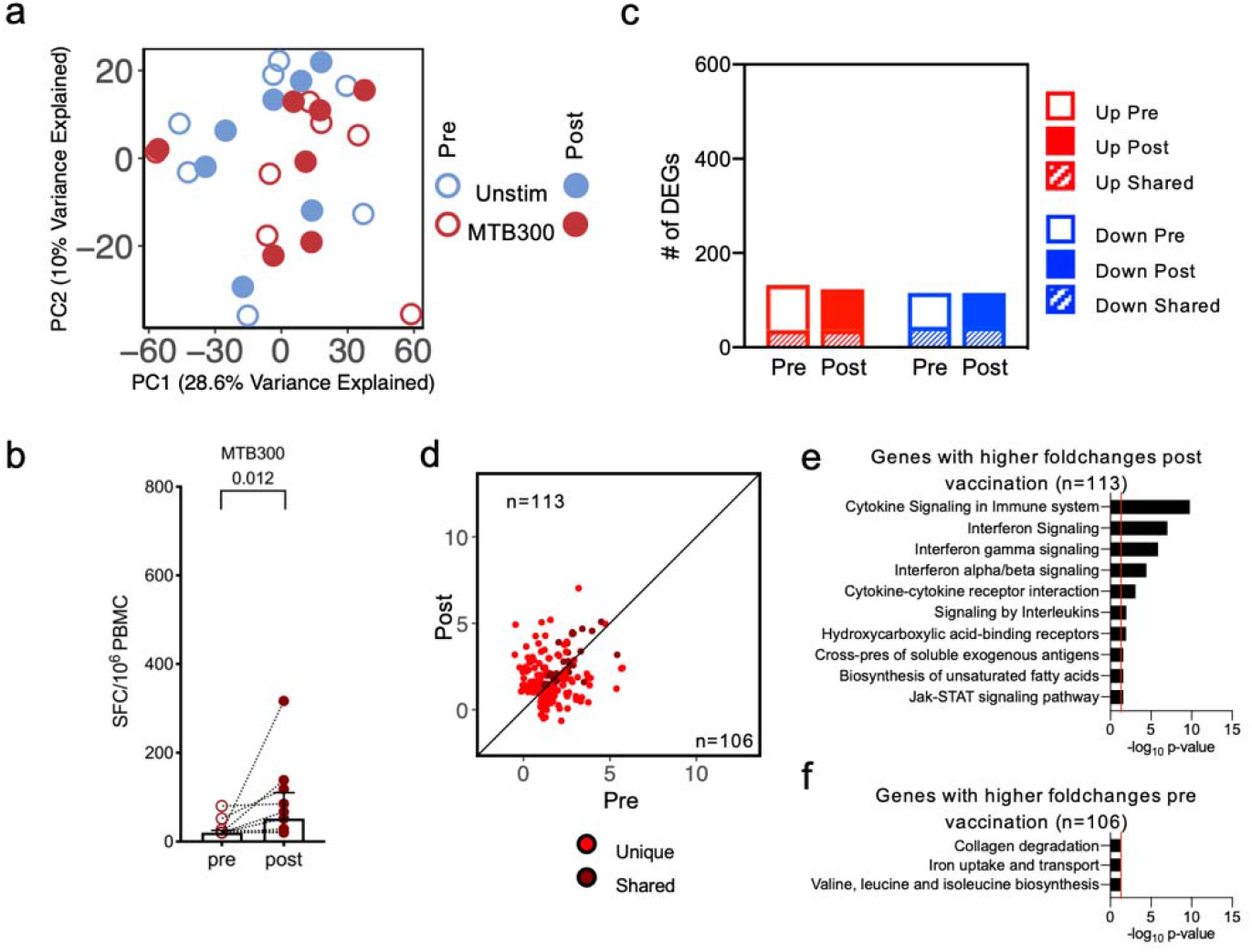
MTB300 peptide pool stimulation has a small impact on BCG vaccination. **a** PCA depicting variation in global gene expression as a result of stimulation condition and vaccination. Empty and filled circles represent the pre- and post-vaccination time points, respectively, and color represents stimulation condition. **b** Paired point graphs depicting magnitude of responses (sum of IFNγ, IL-5, and IL-10) pre- and post-vaccination, against MTB300, as SFC per 10^6^ cultured PBMC. Each point and symbol represent one participant, median ± interquartile range is shown. Wilcoxon matched pair signed rank test. **c** Bar plots representing the total number of differentially expressed genes pre- and post-vaccination. Empty and filled bars represent pre- and post-vaccination, respectively, color represents up or down regulation, and hatched filling represents overlap of genes between the pre and post time points. **d** Scatterplot of genes upregulated upon MTB300 stimulation in pre-vaccination (x-axis) and post-vaccination (y-axis). Each dot represents the log_2_ fold change for a gene, and color represents shared (gene significantly differentially expressed in both pre- and post-vaccination) or unique (differentially expressed in only one condition) expression. Black line at the 45° slope represents identical perturbation in pre- and post-vaccination, with genes above or below the line showing higher perturbation in the post- or pre-group, respectively. **e** Pathway enrichment for genes with higher log_2_ fold changes post vaccination. The ten most significant pathways are shown. **f** Pathway enrichment for genes with higher log_2_ fold changes pre vaccination.

Differential gene expression analysis comparing MTB300 stimulated to unstimulated samples further revealed a lower number of DEGs overall (**Figure 5c; Supplementary table 1**). Additionally, no increase in the number of DEGs was observed post-vaccination (**Figure 5c; Supplementary table 1**). Moreover, there was decreased overlap between pre- and post-vaccination, and comparison of upregulated genes did not show an increased magnitude of log2 fold changes post-vaccination, with distinct sets and similar number of genes pre- and post-vaccination (**Figure 5d; Supplementary table 1**; 113 genes above the 45-degree slope, and 106 genes below the 45-degree slope). Functional enrichment analysis for genes with higher fold changes post-vaccination included pathways such as cytokine and interleukin signaling, similar to that observed in BCG stimulated samples post-vaccination, albeit with lower significance (**Figure 5e; Supplementary table 2)**. There was a high enrichment for interferon signaling upon MTB300 stimulation post-vaccination, with type I, type II and Jak-STAT signaling pathways being significant. In contrast, genes with higher fold changes pre-vaccination were not specifically enriched for any biological functions, with three pathways barely reaching the significance cutoff (**Figure 5f; Supplementary table 2**). These results suggest that, although there is some similarity in the pathways being affected, the MTB300-specific responses are weakly induced following BCG vaccination, especially when compared to the responses upon stimulation with BCG.

### Gene signatures reflect specific cell subset increases post-vaccination

We have previously defined a Th1*-specific gene signature ^25^. Given the likely importance of these Th1* cells in Mtb infection, we further assessed the increase in Th1* cells observed upon BCG vaccination *in silico* in the RNA-sequencing data. Moreover, we have defined a MAIT-specific gene signature ^26^ which was also used *in silico* in the RNA sequencing data. Only upregulated genes from the Th1* and MAIT signatures were used to identify cell-specific gene expression (**Supplementary table 3**). We detected an overall increase in the MAIT and Th1* cell signatures in the BCG stimulated samples compared to the MTB300 stimulated and unstimulated samples (**Supplementary figure 3a and b**). This increase was further enhanced in the BCG post-vaccination samples, similar to the observation in the flow cytometry data. These results suggest an increase in the CD4 T cell frequency following BCG vaccine, specifically in CCR6+ (Th1* and Th17) cell subsets.

### BCG-induced gene expression in PBMCs is primarily driven by CD4 memory T cells

As CD4 T cells are increased upon BCG vaccination, and as memory T cell responses are enhanced upon stimulation with BCG, we also performed RNA-sequencing on CD4 memory T cells isolated from the same individuals as above, pre- and post-vaccination. As with PBMC samples, CD4 memory T cells were stimulated with DMSO (unstimulated), MTB300 peptide pool, and BCG. Differential gene expression analysis comparing BCG stimulated samples to unstimulated samples showed an increase in the number of DEGs post-vaccination (**Figure 6a; Supplementary table 1;** 286 genes upregulated pre-vaccination and 530 genes upregulated post-vaccination, with an overlap of 205 genes), similar to that observed in PBMCs (**Figure 3a**). In contrast, and similar to the observations in PBMCs (**Figure 5c**), MTB300 stimulated samples had a much small number of genes differentially expressed compared to unstimulated samples, and this did not increase post vaccination (**Figure 6a**). Focusing on the effect of BCG stimulation on gene expression, PCA showed a separation between the BCG stimulated and the unstimulated samples across PC1 (**Figure 6b**), however, to a lesser extent than what was observed in PBMCs (**Figure 3a**). Moreover, a reduced separation was observed in pre- and post-vaccination samples within each stimulation condition (**Figure 6b**). Correlation between the unique and shared upregulated DEGs between pre- and post-vaccination BCG stimulated samples showed that the majority of genes had a higher fold change post-vaccination (**Figure 6c; Supplementary table 1;** 467 genes above the 45-degree slope, and 144 genes below the 45-degree slope), also consistent with PBMC analysis (**Figure 3b**). Functional enrichment for genes with higher fold changes post-vaccination identified similar pathways as those observed in PBMC post-vaccination, such as cytokine signaling and interaction and interleukin signaling (**Figure 6d; Supplementary table 2**). Moreover, a Th1*/Th17 type signature was also observed here as evidenced by the IL-23 mediated signaling pathway (**Figure 6d**, showing the 10 most significant pathways), and IL-17 signaling pathway (**Supplementary table 2**). In contrast, the smaller number of genes with higher fold changes pre-vaccination did not result in any significant pathways. The involvement of Th1*/Th17 cells were further strengthened by the observation that there was an increase in BCG-induced IFNγ and IL-17 production in CD4+ T cells following BCG vaccination (**Figure 6e**). No increase in IFNγ production was observed in CD8+ or CD4-CD8-T cells (**Figure 6e**). *In silico* analysis of the Th1* signature (**Supplementary table 3**), as before, showed an increase in the Th1*-specific gene expression in BCG stimulated samples, with a further increase post-vaccination (**Figure 6f**).

**FIGURE 6.**
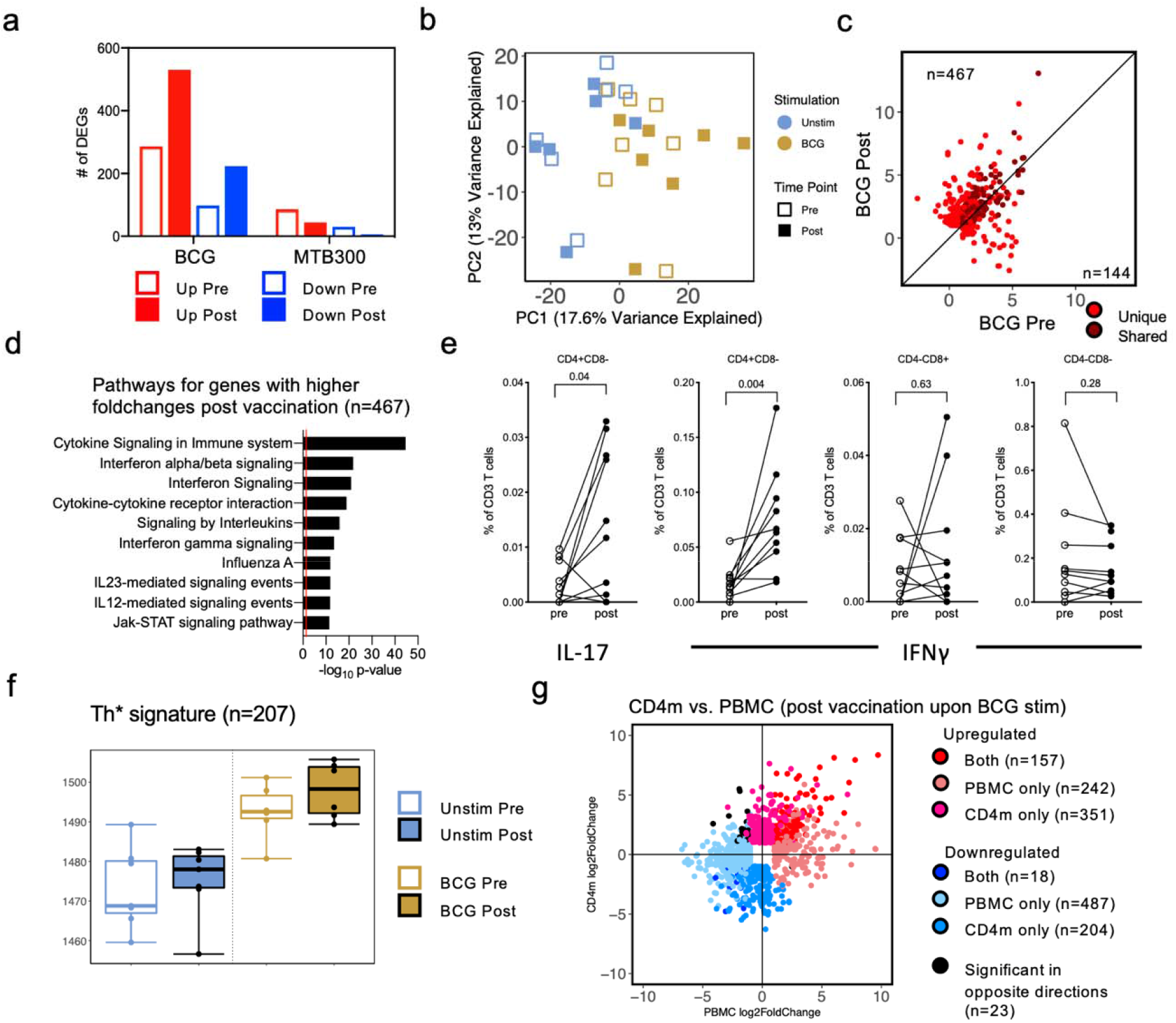
CD4 memory T cells show enhanced gene expression post BCG vaccination in BCG stimulated samples. **a** Grouped bar plots representing differentially expressed genes identified in BCG and MTB300 stimulation, pre- and post-vaccination. Empty bars and filled bars represent pre- and post-vaccination, respectively, and color represents up or down regulation. **b** Principal component analysis depicting the variation in global gene expression as a result of stimulation condition and vaccination. Empty and filled squares represent the pre- and post- vaccination time points, respectively, and color represents stimulation condition. **c** Scatterplot of genes upregulated upon BCG stimulation in pre-vaccination (x-axis) and post-vaccination (y-axis). Each dot represents the log_2_ fold change for a gene, and color represents shared (gene significantly differentially expressed in both pre- and post-vaccination) or unique (differentially expressed in only one condition) expression. Black line at the 45° slope represents identical perturbation in pre- and post-vaccination, with genes above or below the line showing higher perturbation in the post- or pre-group, respectively. **d** Pathway enrichment for genes with higher log_2_ fold changes post vaccination. The ten most significant pathways are shown. **e** Percentage cytokine (IL-17 or IFNγ) detected from T cells in response to BCG. Each dot represents one donor, median ± interquartile range is shown. Wilcoxon matched pair signed rank test. **f** Boxplot of RNA-sequencing data depicting the sum of log_2_ expression values (variance stabilizing transformation, VST) for all genes in the Th1* cell signature. Empty and filled boxplot represent pre- and post-vaccination, respectively, and color represents stimulation condition. Each dot represents an individual donor. **g** Scatterplot of genes differentially expressed between post-vaccination in BCG stimulated compared to unstimulated samples, in PBMC (x-axis) and CD4 memory T cells (y-axis). Each dot represents the log_2_ fold change for a gene, and color represents shared or uniquely differentially expressed genes in PBMC or CD4m.

With the analysis in CD4 memory T cells showing similar results as those observed in PBMCs, we directly compared the DEGs obtained from BCG stimulated compared to unstimulated samples, post-vaccination. Majority of the genes showed similar perturbations in CD4 memory T cells and PBMCs, albeit to differing fold change and significance levels (**Figure 6g; Supplementary table 1**). Only 23 genes were identified to be significant in both PBMC and CD4 memory T cells that had log_2_ fold changes in opposite direction (upregulated in CD4 memory T cells, but downregulated in PBMCs, and vice versa). These results indicate an increase in the magnitude of BCG related gene expression changes post BCG vaccination in CD4 memory T cells. Moreover, the results observed in CD4 memory T cells were similar to those in PBMCs, suggesting that majority of the BCG related gene expression changes observed in PBMCs are driven primarily by CD4 memory T cells, upon BCG vaccination.

### Stimulation with BCG leads to the expansion of specific clonotypes post BCG vaccination

To determine whether BCG vaccination leads to the specific expansion of BCG-reactive clonotypes we performed TCR Sequencing on BCG naïve individuals pre, and 8 months post BCG vaccination. To expand antigen-specific T cells, PBMCs were stimulated with MTB300, a tetanus pool, and BCG for 14 days *in vitro*. Cultures were harvested and DNA was then purified for TCR sequencing using the ImmunoSEQ service from Adaptive Biotechnologies (**Supplementary Table 4**). Every sample had a culture replicate and the ex vivo repertoire of CD4 T cells was utilized as a comparison. We assessed the productive repertoire of each sample, i.e. the unique in-frame rearrangements that do not contain a stop codon, as well as the frequency of these productive clonotypes. We identified a similar number of clonotypes covering eighty percent of the productive repertoire across both replicates across all donor samples in pre- and post-vaccination samples, in all three stimulation conditions (**Figure 7a**). In each stimulated sample there was a selection and expansion of stimuli-specific clonotypes, resulting in fewer unique rearrangements compared to the ex vivo CD4 sample (**Figure 7a**).

**FIGURE 7.**
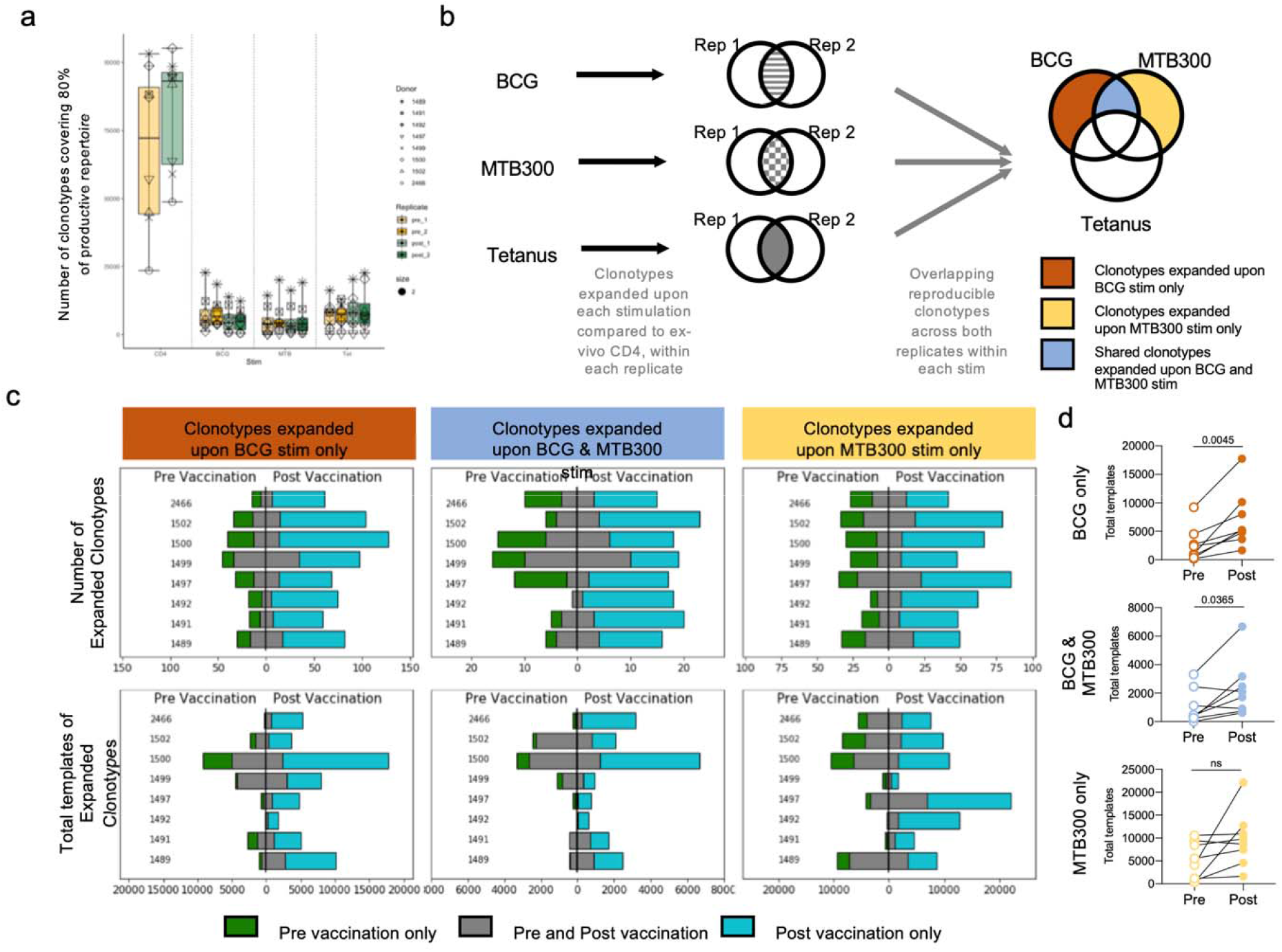
BCG and MTB300 stimulation elicit specific T cell responses. **a** Number of TCR clonotypes covering 80% of the productive repertoire across stimulation conditions and pre- and post-vaccination. Color, yellow and green, represents pre- and post-vaccination, respectively. Each symbol represents one donor. Replicate samples are indicated by the intensity of the color (lighter and darker shade). **b** TCR-sequencing was performed on CD4^+^ T cell samples obtained from BCG naïve individuals (n=8), stimulated with the BCG epitope pool, MTB300 peptide pool, or the tetanus peptide pool. Only those clonotypes that expanded upon stimulation compared to ex-vivo CD4 samples across both replicates were retained within each stimulation condition. The resulting clonotypes in the three stimulation conditions were overlapped to obtain clonotypes that expanded upon BCG stimulation only, MTB300 stimulation only, and clonotypes expanded in both BCG and MTB300 stimulation. Any clonotype expanded upon tetanus stimulation was excluded. This process was repeated for samples obtained post-vaccination. **c** Bar graphs showing the number of expanded clonotypes, and the total templates for these clonotypes, across different conditions, pre- and post-vaccination. Blue indicates clonotypes unique to post-vaccination, green indicates clonotypes unique to pre-vaccination, and grey indicates clonotypes that expanded in both pre- and post-vaccination. **d** Quantification of total templates expanded in pre- and post-vaccination for individual donors in each condition. Empty and filled circles represent pre- and post-vaccination, respectively.

Next, we compared the productive repertoire of samples stimulated with the peptide pools to the corresponding ex vivo CD4 samples, and identified clonotypes that were significantly perturbed as a result of BCG (**Supplementary Figure 4**), and MTB300 and Tetanus stimulation (**Supplementary Figure 5**). A similar number of clonotypes were significantly expanded upon BCG stimulation across all donors, pre-vaccination (**Supplementary Table 5**), and this number was increased post-vaccination (**Supplementary table 5**). To identify reproducible clonotypes, we only retained those clonotypes that expanded significantly across both replicates (-log_2_ OR>1 and FDR p-val<0.05 in both replicates) for an individual donor within each stimulation condition. These reproducible clonotypes within a stimulation condition were overlapped across all three stimulation conditions, to obtain clonotypes that expanded upon BCG stimulation only, MTB300 stimulation only, and clonotypes expanded in both BCG and MTB300 stimulation. Any clonotype expanded upon tetanus stimulation was excluded from further analysis, to remove any non-specific bystander effects associated with the *in vitro* culture (**Figure 7b**). This analysis was performed independently in pre- and post-vaccination samples.

To determine if the clonotypes that expanded post-vaccination corresponded to an increased immune response or simply a result of stimulation, we compared the expanded clonotypes, pre-and post-vaccination, based on the exact clonotype rearrangement sequence. This indicated that although there was an overlap between clonotypes expanded pre- and post-vaccination, this was a very small proportion, and majority of the clonotypes expanded post-vaccination were unique (**Figure 7c**). This suggests that the unique clonotypes that expanded post-vaccination are a result of vaccination, and not stimulation with either the BCG or MTB300 peptide pools. As this overlap was based on the exact clonotype rearrangement sequence, we also performed GLIPH analysis, which clusters TCRs based on similar MHC-restricted peptide antigen binding. The GLIPH analysis revealed a small overlap between pre- and post-vaccination clonotypes (**Supplementary Table 6**), further indicating that the expanded clonotypes post-vaccination are a direct result of the vaccination.

To further examine the homogeneity of TCR repertoires before and after BCG vaccination, we used TCRMatch ^31^ to calculate similarity scores among the CDR3β sequences obtained from the subjects. CDR3β sequences obtained from each subject were separated into four groups depending on vaccination status at the time of sample collection and expansion status in response to BCG stimulation *in vitro*. Scores were computed for pairs of CDR3β sequences within each group, and the fraction of scores exceeding thresholds of 0.84, 0.90, and 0.97 were computed (**Supplementary Figure 6a-c**). The fraction of scores exceeding the threshold was found to be significantly higher in expanded TCRs compared to unexpanded TCRs post-BCG (p=0.016 for 0.84, 0.0029 for 0.90, and p<0.0001 for 0.97). We also found a significant difference between expanded vs. unexpanded TCRs pre-BCG for the lowest threshold (p=0.013 for 0.84) and in the fraction of scores over 0.97 (p=0.0102), which was driven by high similarity scores in one subject. To determine whether repertoire homogeneity was detectable between different subjects, we used TCRMatch to compare CDR3β sequences expanded pre- and post-BCG across subjects (**Supplementary Figure 6d**). The fraction of scores exceeding 0.97 for each cross-subject comparison, separated by BCG vaccination status, revealed that subjects post-BCG had a higher fraction of those scores (14 matches above 0.97 resulting from 9 cross-subject comparisons of post-BCG expanded TCRs, compared to zero matches of pre-BCG expanded TCRs). Of the 14 strong matches between subjects, 7 were found to be exact matches.

We examined the subjects with at least 1 strong match further by assessing similarity between their HLA alleles. All subjects were HLA typed (**Supplementary Table 7**), and therefore the number of shared HLA alleles vs. the fraction of TCRMatch scores above 0.97 could be compared. We found a positive correlation between the number of shared HLA class II alleles and the fraction of TCRMatch scores exceeding 0.97 (**Supplementary Fig. 6e**). The 14 strong matches were also analyzed for evidence of concordance in gene usage. For the 7 identical CDR3β sequences there was an exact match between subjects in all available VDJ and allele calls. For all the strong matches, 13/14 (93%) were determined to have the same V family, that ranged between 2.3-12.4% in prevalence across the subjects.

Finally, the magnitude of expansion, represented by the sum of the number of templates identified for each clonotype (total templates), of the clonotypes pre- and post-vaccination indicated a much greater expansion post-vaccination (**Figure 7c,d**). This significant increase in the degree of expansion was only evident upon stimulation with BCG (in BCG only, and in shared BCG and MTB300), and not upon stimulation with MTB300 only (**Figure 7d**). Taken together, these results indicate both a quantitative and qualitative change in clonotypes post-vaccination. There were new clonotypes observed and an overall expansion of clonotypes post-vaccination upon stimulation with BCG.

## DISCUSSION

Here, we employed a systems biology approach to characterize the T cell subsets, cytokine secretory profiles, and other cell subset markers that change following BCG vaccination to yield insight into potential new correlates of protection. Specifically, total PBMC and purified CD4+ T cell population transcriptomics were used in combination with DNA methylome analysis and other immunological techniques, such as flow cytometry and fluorospot, to characterize BCG-induced responses in a longitudinal adult cohort who provided blood samples 1-2 weeks prior to and 8 months following BCG vaccination. Gaps in knowledge surrounding BCG-induced immunity, both in terms of antigen-specificity and functional responses, particularly the part of the response associated with protection against *Mtb* that should be boosted by a vaccine ^32-34^, impede efforts to improve variable BCG efficacy. Few studies have used multi-omics approaches in the context of BCG vaccination and were conducted mostly in animals. Cortes et al. conducted a transcriptomic analysis in mice comparing BCG vaccinated and naïve mice before and after *M. bovis* challenge and found Th17-associated cytokines correlated with protection ^35^. Darrah et al. used a combination of immunohistochemistry, flow cytometry and single cell sequencing to characterize the immune response in macaques after intravenous administration of BCG, a route that was largely successful as 9/10 vaccinated animals were protected even after challenge with virulent *Mtb* 6 months post vaccination ^4^. Similarly, Hoft et al. used systems immunology to show that oral and mucosal BCG delivery induced distinct molecular signatures, which could potentially permit the identification of genes that should be differentially targeted by vaccines geared toward inducing optimal systemic or mucosal TB immunity ^36^. A transcriptional signature for BCG will serve as an important comparator for novel vaccination strategies, facilitating their design and evaluation.

Our results reaffirm the critical role of T cells, particularly CD4 T cells, in mediating antimycobacterial immunity ^37^, both in the context of *Mtb* infection and vaccination ^20,38-40^. The response at the mRNA level observed from post-BCG CD4 memory T cells was very similar to those in post-BCG PBMCs, suggesting that the majority of the BCG-induced gene expression changes observed in PBMCs are driven primarily by CD4 memory T cells upon BCG vaccination. Other studies have found that BCG induces higher CD4 T cell responses than CD8 responses ^22,41^, and with increased levels of BCG-reactive cells post-vaccination ^21,22^.

We further characterized the particular cell subset within the CD4 compartment that mediates BCG-induced responses through cell surface phenotypes and gene signatures and found that it is associated with Th1* (CXCR3+CCR6+CCR4-) cells. Our group previously showed that this CD4+ Th subset contained the majority of *Mtb*- ^42^ and NTM-specific T cells ^27^. Moreover, we later showed that Th1* was involved in the immune response following natural *Mtb* infection, since Th1* was increased in individuals with LTBI compared to TB negative controls ^25^. Here, we found an increase of CCR6+ CD4 T cells (including Th1* and Th17 subsets) and CD3+MR1+ MAITs following BCG vaccination. These frequency changes could also be recapitulated *in silico* in the RNA-seq data as detected by an overall increase in previously characterized MAIT and Th1* cell subset signatures ^25,26^ post vaccination. The increase in the frequency of MAIT cells observed here is in contrast to what was recently found in infants following primary BCG vaccination and following BCG revaccination in tuberculin skin test positive adults (consistent with prior *Mtb* infection) ^43^, which are both different from our cohort of primary vaccinated adults. Providing further evidence for the importance of the CCR6+ CD4 T cells, we also found increases in IL-23 mediated and IL-17 signaling pathways following BCG stimulation post vaccination both in PBMC and CD4 memory T cells. In addition, we found an increase in the magnitude of BCG-induced CCR6+ associated IFNγ and IL-17 production in CD4+ T cells following BCG vaccination.

Several studies suggest an important role for these antigen-specific Th1* cells in the immune response against mycobacteria and they may be a promising candidate for a correlate of protection against *Mtb*. Th1* cells also share characteristics with a CCR6+ CD4 cell subset recently described as preferentially enriched in a cohort of TB non-progressors compared to those who progressed to active TB ^44^. In non-human primates (NHP), antigen-specific CD4 T cells in the PBMCs and bronchoalveolar lavage fluid of rhesus macaques that received BCG intravenously had a similar Th1/Th17 phenotype and importantly the majority of these animals (9/10) were protected from *Mtb* challenge 6 months post-BCG ^4^. This Th1*-like cell subset has also been shown to be associated with protection against *Mtb* in other NHP studies ^45,46^. Given the high diversity in results and outcomes across studies and models, a correlate of vaccine-induced protection will most likely not be a single marker, which makes it important to study additional cell markers such as activation, migration and memory markers, as well as functional secretory profiles in order to separate immunopathology from protective antigen-specific T cell responses that can serve as correlates of protection ^47-49^.

In the present study, we also found ligands for CXCR3, CXCL10 (IP-10) and CXCL11 that exhibited a higher fold-change post-vaccination. Antigen-specific Th1* CD4 T cells express the tissue homing chemokine receptor CXCR3 and could thus respond to these ligands. It has been proposed that post-vaccination measurement of multifunctional responses in *Mtb*-specific, relatively undifferentiated, memory T cell subsets retaining the capacity to traffic to the lung may be more indicative of protective immunity against TB ^50^. Murine models have implicated migration markers, including CXCR3 ^51,52^, that confer the ability of CD4 T cells to exit the circulation and enter the lung to interact with *Mtb*-infected APCs, as promising correlates of protection candidates.

Reactivity against our *Mtb*-derived peptide pool defined in healthy *Mtb* infected interferon gamma release assay (IGRA) positive individuals, MTB300, was also boosted post-BCG vaccination, however to a lesser degree than BCG-induced immune responses. MTB300 contains 255 peptides out of a total of 300 that are homologous between *Mtb* and BCG. This result provides further evidence that the antigen-specific human T cell responses triggered by BCG vaccination are not fully understood. Determination of BCG-specific epitopes and antigens will provide crucial immune monitoring reagents.

Most genes upregulated post-BCG vaccination also tend to be upregulated, albeit to a lesser extent, pre-vaccination. This suggests a boosting of the response that is present pre-vaccination. The BCG-specific response is probably heavily influenced by exposure to mycobacteria that the immune system has been primed with in the past. Moreover, BCG vaccination resulted in an overall increase in the BCG-induced response, but DEGs and BCG-specific immune responses were also identified pre-vaccination following BCG stimulation. This can be, at least partly, explained by previous NTM exposure resulting in cross-reactive immune responses ^27^. The signatures we have identified here can be compared to those found in children that were vaccinated at birth and following BCG revaccination, when available. Especially given the protection from severe TB observed in children and lack of efficacy in adults mediated by BCG.

BCG vaccination resulted in expanded TCR repertoires with higher intra-repertoire homogeneity than unexpanded TCRs, as well as similar TCR repertoires across different subjects. Thus, the TCRs appear to converge towards similar TCR sequences more than they diversify following BCG vaccination. Clustered TCRs have been described previously in the responses to herpesviruses and *Mtb* ^53,54^. These similar TCR repertoires may be influenced by similarities in HLA alleles. Future studies can reveal how certain HLA alleles influence BCG vaccine efficacy and whether this can be used as a means of predicting efficacy.

In conclusion, this study provides a detailed characterization of BCG-induced immune responses and TCR clonotypes in adults that were vaccinated with BCG. These findings inform our understanding of the immune response induced by the BCG vaccine and provides means to track longitudinal changes in the specific T cells in many different settings.

## MATERIALS AND METHODS

### Ethics statement

All participants provided written informed consent for participation in the study. Ethical approval was obtained from the Institutional review boards at Linköping University (2015/150-32) and the La Jolla Institute for Immunology (VD-140).

### Study subjects

We recruited 17 TB negative and BCG naïve individuals for participation in the study who were offered the BCG vaccine through their medical school at Linköping University. Their *Mtb* infection status was confirmed by a negative tuberculin skin test and IFNγ-release assay (IGRA; T.Spot-TB, Oxford Immunotec).

Venous blood was collected in heparin-containing blood bags 1-2 weeks prior to BCG vaccination and 8 months after the vaccination. Peripheral blood mononuclear cells (PBMC) were purified from whole blood by density-gradient centrifugation (Ficoll-Hypaque, Amersham Biosciences), according to the manufacturer’s instructions. Cells were cryopreserved in liquid nitrogen suspended in FBS (Gemini Bio-Products) containing 10% (vol/vol) DMSO (Sigma). Cryopreserved cells were shipped from Linköping University to LJI for analysis.

### HLA typing

Participants were HLA typed by an ASHI-accredited laboratory at Murdoch University (Institute for Immunology & Infectious Diseases, Western Australia) as previously described ^55^. HLA typing for class I (HLA A, B, C) and class II (DQA1, DQB1, DRB1, 3, 4, 5, DPB1) was performed using locus-specific PCR amplification of genomic DNA. Patient-specific, barcoded primers were used for amplification. Amplified products were quantitated and pooled by subject and up to 48 subjects were pooled. An indexed (8 indexed MiSeq runs) library was then quantitated using Kappa universal QPCR library quantification kits. Sequencing was performed using an Illumina MiSeq using 2×300 paired-end chemistry. Reads were quality-filtered and passed through a proprietary allele calling algorithm and analysis pipeline using the latest IMGT HLA allele database as a reference. The algorithm was developed by E.J.P. and S.A.M. and relies on periodically updated versions of the freely available international immunogenetics information system (http://www.imgt.org) and an ASHI-accredited HLA allele caller software pipeline, IIID HLA analysis suite (http://www.iiid.com.au/laboratory-testing/). The HLA type of each subject is listed in **Supplementary table 7**.

### Peptides and other stimuli

Peptides were synthesized as crude material on a small (1mg) scale by A&A, LLC (San Diego, CA). Multi-epitope peptide pools (“megapools”) were prepared as previously described ^56^. Individual peptides were resuspended in DMSO, and equal amounts of each peptide were pooled to contruct the peptide pool. After lyophilization, the peptide pools were resuspended in DMSO, aliquoted, and stored at -20°C. Two different peptide pools were used; a peptide pool containing 300 Mtb-derived 15-20-mer peptides (MTB300) primarily HLA class II restricted ^56^, and a peptide pool with 125 peptides derived from the *Clostridium tetani* toxin (TT) sequence ^57^.

In addition to peptide pool, PBMCs were also stimulated with *M. bovis* BCG-Danish or Pasteur at 100 µg/ml.

### Fluorospot assay

Antigen-specific cellular responses were measured by IFNγ Fluorospot assay with all antibodies and reagents from Mabtech (Nacka Strand, Sweden). Plates were coated overnight at 4°C with a mouse anti-human IFNγ (clone 1-D1K) antibody. Briefly, 2×10^5^ cells were added to each well of pre-coated Immobilon-FL PVDF 96-well plates (Mabtech) in the presence of 2 μg/ml peptide pool, or 100µh/ml BCG, and incubated at 37°C in humidified CO_2_ incubator for 20-24 hrs. Cells stimulated with DMSO (corresponding to the percent DMSO in the peptide pools) were used to assess non-specific/background cytokine production and PHA stimulation at 10 μg/ml was used as a positive control. All conditions were tested in triplicates. Fluorospot plates were developed according to manufacturer’s instructions (Mabtech). Briefly, cells were removed and plates were washed 6 times with 200 μl PBS/0.05% Tween 20 using an automated plate washer. After washing, 100 μl of antibody mixture containing anti-IFNγ (7-B6-1-FS-FITC) prepared in PBS with 0.1% BSA was added to each well and plates were incubated for 2 hrs at room temperature. Plates were again washed 6 times with 200 μl PBS/0.05% Tween 20 using an automated plate washer and incubated with diluted fluorophores (anti-BAM-490) for 1 hr at room temperature. Finally, plates were once more washed 6 times with 200 μl PBS/0.05% Tween 20 using an automated plate washer and incubated with fluorescence enhancer for 15 mins at room temperature. The plates were blotted dry and spots were counted by computer-assisted image analysis (AID iSpot, Aid Diagnostica GMBH, Strassberg, Germany). Responses were considered positive if the net spot-forming cells (SFC) per 10^6^ PBMC were ≥20, the stimulation index ≥2, and p≤0.05 by Student’s t-test or Poisson distribution test.

### Flow cytometry

Several different flow cytometry panels were used. Cryopreserved PBMCs were thawed in RPMI supplemented with 5% human serum (Gemini Bio-Products, West Sacramento, CA), 1% Glutamax (Gibco,Waltham,MA), 1% penicillin/streptomycin (Omega Scientific, Tarzana, CA), and 50U/ml Benzonase (Millipore Sigma, Burlington, MA). Cells were then washed and counted. 1 million cells were then blocked in 10% FBS for 10 mins at 4°C. After blocking, cells were stained with APCef780 conjugated anti-CD4 (clone RPA-T4, eBiosciences), AF700 conjugated anti-CD3 (UCHT1, BD Pharmigen), BV650 conjugated anti-CD8a (RPA-T8, Biolegend), PECy7 conjugated anti-CD19 (HIB19, TONBO), APC conjugated anti-CD14 (61D3, TONBO), PerCPCy5.5 conjugated anti-CCR7 (G043H7, Biolegend), PE conjugated anti-CD56 (CMSSB, eBiosciences), FITC conjugated anti-CD25 (M-A251, BD Pharmigen), eF450 conjugated anti-CD45RA (HI100, eBiosciences) and fixable viability dye eF506 (eBiosciences) for 30 mins at 4°C. Cells were then washed twice and acquired on a BD FACSAria flow cytometer (BD Biosciences, San Jose, CA) to measure the frequency of different cell subsets. The gating strategy for this panel was performed as previously reported ^24^.

Cells were also stained with BV650 conjugated CCR6 (G034E3, BioLegend), CXCR3-APC (1C6/CXCR3, BD Biosciences) for 20 min at 37°C, followed by CCR4-PE-Cy7 (1G1, BDBiosciences), CCR7-PerCPCy5.5 (UCHL1, BioLegend), CD4-APCef780 (RPA-T4, eBiosciences), CD3-AF700 (UCHT1, BD Pharmigen) CD45RA-eF450 (HI100, eBiosciences), CD8-V500 (RPA-T8, BD Biosciences), CD14-V500 (M5E2; BD Biosciences), CD19-V500 (HIB19, BD Biosciences), and fixable viability dye eF506 (eBiosciences) at room temperature for 30 min. Gating strategy is shown in **Supplementary Figure 7a**.

For non-conventional T cells PBMCs were stained with 1:100 MR1 5-OP-RU or 6-FP (as a control) tetramer for 40 min at room temperature. The MR1 tetramer technology was developed jointly by Dr. J. McCluskey, Dr. J. Rossjohn, and Dr. D. Fairlie ^58^, and the material was produced by the National Institutes of Health Tetramer Core Facility, as permitted to be distributed by the University of Melbourne. After 40 min, cells were also stained with fixable viability dye eF506 (eBiosciences) and with CD3-AF700 (UCHT1, BD Pharmigen), CD4-APCef780 (RPA-T4, eBiosciences), CD8-BV650 (RPA-T8; BioLegend), CD14-V500 (M5E2; BD Biosciences), CD19-V500 (HIB19, BD Biosciences), CD161-APC (HP-3G10; eBiosciences), Vα7.2-PE-Cy7 (3C10, BioLegend), Vα24-PE-Dazzle594 (6B11, BioLegend), and γδPAN TCR-FITC (11F2, BD Biosciences) for 30 min at room temperature. The gating strategy is shown in **Supplementary Figure 7b**. MR1+ T cells were defined as 5-OP-RU MR1 tetramer+, and CD4-, Vα24-, γδPAN TCR-, Vα7.2+, and CD161+ through Boolean gating.

For the flow cytometry measurement of IFNγ and IL-17, PBMCs were thawed and stimulated with 100µg/ml BCG or left unstimulated in the presence of 1µg/ml anti-CD28 (CD28.2 eBioscience), and 1µg/ml anti-CD49d (9F10, BioLegend). Cells were incubated at 37°C for 5 hours, after which 2.5µg/ml Brefeldin A and monensin was added for another 7 hours. Cells were washed and blocked in 10% FBS for 10 mins at 4°C. Cells were then stained with CD4-APCef780 (RPA-T4, eBiosciences), CD3-AF700 (UCHT1, BD Pharmigen), CD8-BV650 (RPA-T8; BioLegend), CD14-V500 (M5E2; BD Biosciences), CD19-V500 (HIB19, BD Biosciences), fixable viability dye eF506 (eBiosciences) for 30 min at 4°C. Cells were washed twice and then fixed in 4% paraformaldehyde solution for 10 min at 4°C. Saponin buffer was used to permeabilize the cells by incubating them at room temperature for 10 min, followed by blocking in 10% FBS for 5 min at 4°C. Cells were then stained with IFNγ-FITC (4S.B3, eBioscience) and IL-17-PE-Cy7 (eBio64DEC17, eBioscience) for 30 min at room temperature. Followed by washes and acquisition. The gating strategy is shown in **Supplemental figure 7c**.

### Fluorescence-activated cell sorting

PBMCs were thawed and 2×10^6^ cells were added per well in a 96 round bottom well plate. Cells were stimulated with BCG (50µg/ml), MTB300 (2µg/ml), Tetanus pool (2µg/ml), or DMSO (corresponding to the percent DMSO in the peptide pools) as a control. Anti-human CD28 and CD3 (1 µg/ml) was used as a positive control. The wells for the positive control was pre-coated overnight at 4°C. Cells were incubated at 37°C for 24 h.

The following day, cells were washed and incubated in PBS with 10% FBS at 4°C for 10 min. They were stained with fixable viability dye eFluor 506 (eBioscience), and an antibody cocktail containing anti-human CD3-Alexa Fluor 700 (UCHT1, BD Bioscience), CD4-APCeFluor 780 (RPA-T4, eBioscience), CD8-V500 (RPA-T8, BD Biosciences), CD45RA-eFluor 450 (HI100, eBioscience), and CCR7-PerCPCy5.5 (UCHL1, BioLegend) for 20 min at room temperature. Cells were transferred into a 5 ml polypropylene FACS tube (BD Biosciences) and PBMCs (excluding doublets) and CD4 memory T cells were sorted on a FACSAria III cell sorter (Becton Dickinson) into QIAzol Lysis Reagent (QIAGEN). A total of 100,000 cells was sorted per sample. For gating strategy see **Supplementary figure 7d**. Sorted cell populations were stored in QIAzol Lysis Reagent at -80°C until RNA extraction.

### CD4 T cell isolation

CD4+ T cells were isolated from at least 5×10^6^ PBMCs on the day of thaw by negative selection using the CD4+ T cell isolation kit II (Miltenyi Biotec, Bergisch Gladbach, Germany) according to manufacturer’s instructions. The isolated CD4+ T cells were washed, pelleted, and then stored at -80°C until DNA extraction.

### Cell expansion for TCR sequencing

For *in vitro* expansion, cryopreserved PBMCs were thawed in RPMI supplemented with 5% human serum (Gemini Bio-Products, West Sacramento, CA), 1% Glutamax (Gibco, Waltham, MA), 1% penicillin/streptomycin (Omega Scientific, Tarzana, CA), and 50 U/ml Benzonase (Millipore Sigma, Burlington, MA). The cells were then washed and viability was evaluated using trypan blue dye exclusion. Briefly, at a density of 2×10^6^ cells per mL, the cells were plated in a well of a 24-well plate in the presence of 50µg/ml BCG, 2µg/ml MTB300, or 2µg/ml tetanus peptide pool, and were incubated in a 37°C humidified CO_2_ incubator for 14 days. Every 3-4 days cells were supplied with 10 U/ml recombinant human IL-2. After 14 days of culture, cells were harvested, counted, and pelleted. The cell pellets were stored at -80°C until DNA extraction.

### RNA sequencing

RNA sequencing was performed as described previously (24). Briefly, total RNA was purified using an miRNeasy Micro Kit (QIAGEN) and quantified by quantitative PCR, as described previously (25). Purified total RNA (1–5ng) was amplified following the Smart-Seq2 protocol (16 cycles of cDNA amplification) (26). cDNA was purified using AMPure XP beads (Beckman Coulter). From this step, 1 ng of cDNA was used to prepare a standard Nextera XT sequencing library (Nextera XT DNA sample preparation kit and index kit, Illumina). Whole-transcriptome amplification and sequencing library preparations were performed in a 96-well format to reduce assay-to-assay variability. Quality-control steps were included to determine total RNA quality and quantity, the optimal number of PCR preamplification cycles, and fragment size selection. Samples that failed quality control were eliminated from further downstream steps. Barcoded Illumina sequencing libraries (Nextera; Illumina) were generated using the automated platform (Biomek FXp). Libraries were sequenced on a NovaSeq 6000 Illumina platform to obtain 50-bp paired end reads (TruSeq Rapid kit; Illumina).

### TCR sequencing

DNA was extracted from the cultured cells or ex vivo CD4^+^ T cell samples using DNeasy Blood and Tissue kit (Qiagen, Hilden, Germany) according to manufacturer’s instructions. Samples were sent to Adaptive Biotechnologies (Seattle, WA) for TCRB sequencing according to their protocol. The ex vivo CD4^+^ T cell samples were sequenced with “deep resolution” to cover a maximum number of clonotypes in the repertoire. Samples that were stimulated with peptide pools for 14 days and then harvested were sequenced with “survey resolution”.

### Data analysis – RNA sequencing

Paired-end reads that passed Illumina filters were filtered for reads aligning to tRNA, rRNA, adapter sequences, and spike-in controls. The reads were aligned to the GRCh38 reference genome and Gencode v27 annotations using TopHat v1.4.1 ^59^. DUST scores were calculated with PRINSEQ Lite v0.20.3 ^60^ and low-complexity reads (DUST > 4) were removed from BAM files. The alignment results were parsed via SAMtools ^61^ to generate SAM files. Read counts to each genomic feature were obtained with the htseq-count program v0.7.1 ^62^ using the “union” option. Raw counts were imported into R v3.6.1 where they were subset into PBMC and CD4 memory, both containing unstimulated, MTB300 peptide pool, and BCG epitope pool stimulated samples. The following steps were performed on PBMC and CD4 memory subsets independently. R/Bioconductor package DESeq2 v.1.24.0 ^63^ was used to normalize raw counts. Variance stabilizing transformation was applied to normalized counts to obtain log_2_ gene expression values. Quality control was performed using boxplots and Principal component analysis (PCA), using the ‘prcomp’ function in R, on log_2_ expression values. Differentially expressed genes were identified using the DESeq2 Wald test, and p-values were adjusted for multiple test correction using the Benjamini Hochberg algorithm ^64^ (**Supplementary Table 1**). Genes with adjusted p values < 0.05 and log2 fold change > 1 or < -1 were considered differentially expressed. Pathway enrichment analysis was performed using ToppGene Suite: ToppFun ^65^ (**Supplementary Table 2**), and cell type enrichment was performed using DICE ^66^.

### Data analysis – TCR sequencing

Pre-processing and quality control of the raw data was performed using the immunoSEQ analyzer (Adaptive Biotechnologies, Inc.). Measurement metrics of processed data were exported in the tsv file format and downstream data analysis was performed in Python v3.7.2 and in R v3.6.1. To identify clonotypes that were expanded in culture or after vaccine, each of the replicates per donor was compared to the corresponding ex vivo CD4 sample, and p-values and odds-ratio were calculated using a two-sided Fisher exact test, using the ‘fisher_exact’ function in the SciPy.Stats v1.4.1 ^67^ and NumPy v1.17.2 ^68^ extensions of python. Clonotypes appearing in both replicates with -log_2_ odds ratios (OR) >1 or <-1 and false discovery rate (FDR) p-value□ < □0.05 corrected for multiple testing using the Benjamini–Hochberg method ^64^, calculated using the ‘fdrcorrection’ function from the statsmodels module v0.9.0 ^69^ for Python, were considered significant (**Supplementary Table 5**). For visualization purposes, all -log_10_ FDR p-values > 50 were set to 50. Sequence similarity by clustering was performed using GLIPH v1.0 with default parameters, to identify conserved motifs and global similarity of complementarity-determining region 3 (CDR3) sequences ^53^ (**Supplementary Table 6**).

We also used TCRMatch to determine sequence similarity ^31^. Within each group, defined by a single subject, vaccination and expansion status, and CDR3β sequences were tested against each other. For groups with expanded sequences, all CDR3β were analyzed against each other. For groups with unexpanded TCRs, the number of CDR3β sequences were much greater than the number of expanded CDR3β sequences in the same individual, so random sampling was performed. A sample of n sequences, with n=the number of expanded CDR3β sequences for the corresponding individual and vaccination status, was randomly selected and subsequently run through TCRMatch to assess intragroup similarity. 100 random samplings were performed for each group of unexpanded sequences. Cross-subject comparisons were performed by running TCRMatch on all possible pairs of sequences between all possible pairs of subjects. The number of HLA alleles shared by two subjects was calculated by comparing each subjects’ HLA alleles for a given HLA locus against another subjects’ corresponding HLA alleles. All pairwise combinations were compared for each gene, with a maximum of four shared allele pair combinations. For example, if subject 1 carried alleles X and Y, and subject 2 carried X and Z, the number of shared alleles was recorded as 1. Meanwhile, if subject 1 were homozygous for X, and subject 2 carried X and Z, the number of shared alleles was recorded as 2. Finally, if both subjects were homozygous for the same allele, their number of shared allele pairs was counted as 4.

### DNA methylome data acquisition and analyses

The DNA methylome data was analyzed using the HumanMethylationEPIC (850K) array (Illumina, USA) as per manufacturer’s instruction. The raw IDAT files of the DNA methylome data was processed using the ChAMP package ^70^ in R (v4.0.3) after using the default filtering criteria i) removing the CpGs with detection p-value >0.01, ii) filter out CpGs with <3 beads in at least 5% of samples per CpGs, iii) filter out all non-CpGs contained in the dataset, iv) removing all SNP-related CpGs, v) filter out all multi-hit probes, and vi) filter out all CpGs located in X and Y chromosomes. The filtered data was normalized using the β mixture quantile normalization (BMIQ) function using the ChAMP package. For each CpG site, the methylation β values were calculated, which represents the fraction of methylated cytosines at their particular CpG site (0=unmethylated, 1=fully methylated). The Houseman algorithm ^30^ was used to calculate the cell type deconvolution as the samples were drawn from the PBMCs for “before” and “after” samples separately. We performed a Shapiro-Wilk test to test the normality of the samples and used a t-test to compare differences between the before and after group for each cell type. The differential methylation values (mean methylation difference, mmd) were calculated using the two different groups of samples, “before” and “after”. A total of 139 613 CpGs were identified as differentially methylated CpGs (DMCs). A volcano plot was generated using the EnhancedVolcano package of hypermethylated and hypomethylated CpGs. The differentially methylated CpGs were annotated using the human genome annotation (HG38.13) to find the corresponding genes (Differentially Methylated Genes, DMGs). The hyper and hypomethylated CpGs were annotated with the different chromosomal locations. The hyper and hypomethylated genes were compared with the up and downregulated differentially expressed genes using the venneuler package ^71^ (in house python script). A principal component analysis (PCA) was performed on the normalized β values of DNA methylome dataset using the factoMineR and factoExtra (https://CRAN.R-project.org/package=factoextra) packages. All DMCs were considered significant with the Benjamini-Hochberg (BH) corrected p-value < 0.15, if not stated otherwise. A list of DMGs intersected with DEGs were analyzed using the clusterProfiler package ^72^ using the Kyoto Encyclopedia of Genes and Genomes (KEGG) database.

## Data availability

The RNA-seq datasets analyzed as part of this study have been deposited in the NCBI Gene Expression Omnibus (GEO) database with the primary accession number GSE156422.

## AUTHOR CONTRIBUTIONS

ASi, PD, ASe, BP, and CSLA participated in the design and direction of the study. ASi, PD, RK, KM, WC, JD, ML, and CSLA performed and analyzed experiments. GS and PV performed the RNAseq. PD performed the bioinformatics analysis, supervised by ASi. ML recruited participants and maintained participant data. EJP and SAM coordinated and performed HLA typing. ASi, PD, KM, ML, and CSLA wrote the manuscript. All authors read, edited and approved the manuscript.

## ACKNOWLEDGMENTS

This study was supported by U19 AI118626, S10 RR027366, and S10 OD016262.

## REFERENCES

1 Colditz, G. A. et al. The efficacy of bacillus Calmette-Guerin vaccination of newborns and infants in the prevention of tuberculosis: meta-analyses of the published literature. Pediatrics 96, 29–35 (1995).

2 Fine, P. E. Variation in protection by BCG: implications of and for heterologous immunity. Lancet 346, 1339–1345 (1995).

3 Trunz, B. B., Fine, P. & Dye, C. Effect of BCG vaccination on childhood tuberculous meningitis and miliary tuberculosis worldwide: a meta-analysis and assessment of cost-effectiveness. Lancet 367, 1173–1180, doi:10.1016/S0140-6736(06)68507-3 (2006).

4 Darrah, P. A. et al. Prevention of tuberculosis in macaques after intravenous BCG immunization. Nature 577, 95–102, doi:10.1038/s41586-019-1817-8 (2020).

5 Manjaly Thomas, Z. R. & McShane, H. Aerosol immunisation for TB: matching route of vaccination to route of infection. Transactions of the Royal Society of Tropical Medicine and Hygiene 109, 175–181, doi:10.1093/trstmh/tru206 (2015).

6 White, A. D. et al. Evaluation of the Immunogenicity of Mycobacterium bovis BCG Delivered by Aerosol to the Lungs of Macaques. Clinical and vaccine immunology : CVI 22, 992–1003, doi:10.1128/CVI.00289-15 (2015).

7 Price, D. N., Kusewitt, D. F., Lino, C. A., McBride, A. A. & Muttil, P. Oral Tolerance to Environmental Mycobacteria Interferes with Intradermal, but Not Pulmonary, Immunization against Tuberculosis. PLoS Pathog 12, e1005614, doi:10.1371/journal.ppat.1005614 (2016).

8 Barreto, M. L. et al. Causes of variation in BCG vaccine efficacy: examining evidence from the BCG REVAC cluster randomized trial to explore the masking and the blocking hypotheses. Vaccine 32, 3759–3764, doi:10.1016/j.vaccine.2014.05.042 (2014).

9 Dourado, I. et al. Rates of adverse reactions to first and second doses of BCG vaccination: results of a large community trial in Brazilian schoolchildren. Int J Tuberc Lung Dis 7, 399–402 (2003).

10 Rodrigues, L. C. et al. Effect of BCG revaccination on incidence of tuberculosis in school-aged children in Brazil: the BCG-REVAC cluster-randomised trial. Lancet 366, 1290–1295, doi:10.1016/S0140-6736(05)67145-0 (2005).

11 Bekker, L. G. et al. A phase 1b randomized study of the safety and immunological responses to vaccination with H4:IC31, H56:IC31, and BCG revaccination in Mycobacterium tuberculosis-uninfected adolescents in Cape Town, South Africa. EClinicalMedicine 21, 100313, doi:10.1016/j.eclinm.2020.100313 (2020).

12 Nemes, E. et al. Prevention of M. tuberculosis Infection with H4:IC31 Vaccine or BCG Revaccination. N Engl J Med 379, 138–149, doi:10.1056/NEJMoa1714021 (2018).

13 Rakshit, S. et al. BCG revaccination boosts adaptive polyfunctional Th1/Th17 and innate effectors in IGRA+ and IGRA-Indian adults. JCI Insight 4, doi:10.1172/jci.insight.130540 (2019).

14 Nieuwenhuizen, N. E. & Kaufmann, S. H. E. Next-Generation Vaccines Based on Bacille Calmette-Guerin. Front Immunol 9, 121, doi:10.3389/fimmu.2018.00121 (2018).

15 Scriba, T. J. et al. Dose-finding study of the novel tuberculosis vaccine, MVA85A, in healthy BCG-vaccinated infants. The Journal of infectious diseases 203, 1832–1843, doi:10.1093/infdis/jir195 (2011).

16 McShane, H. et al. Recombinant modified vaccinia virus Ankara expressing antigen 85A boosts BCG-primed and naturally acquired antimycobacterial immunity in humans. Nature medicine 10, 1240–1244, doi:10.1038/nm1128 (2004).

17 Nieuwenhuizen, N. E. et al. The Recombinant Bacille Calmette-Guerin Vaccine VPM1002: Ready for Clinical Efficacy Testing. Front Immunol 8, 1147, doi:10.3389/fimmu.2017.01147 (2017).

18 Abebe, F. Is interferon-gamma the right marker for bacille Calmette–Guérin-induced immune protection? The missing link in our understanding of tuberculosis immunology. Clinical & Experimental Immunology 169, 213–219, doi:10.1111/j.1365-2249.2012.04614.x (2012).

19 Kagina, B. M. et al. Specific T cell frequency and cytokine expression profile do not correlate with protection against tuberculosis after bacillus Calmette-Guerin vaccination of newborns. American journal of respiratory and critical care medicine 182, 1073–1079, doi:201003-0334OC [pii] 10.1164/rccm.201003-0334OC (2010).

20 Fletcher, H. A. et al. T-cell activation is an immune correlate of risk in BCG vaccinated infants. Nature communications 7, 11290, doi:10.1038/ncomms11290 (2016).

21 Soares, A. P. et al. Bacillus Calmette-Guerin Vaccination of Human Newborns Induces T Cells with Complex Cytokine and Phenotypic Profiles. J Immunol 180, 3569–3577 (2008).

22 Rodo, M. J. et al. A comparison of antigen-specific T cell responses induced by six novel tuberculosis vaccine candidates. PLoS Pathog 15, e1007643, doi:10.1371/journal.ppat.1007643 (2019).

23 Burel, J. G. et al. Host Transcriptomics as a Tool to Identify Diagnostic and Mechanistic Immune Signatures of Tuberculosis. Front Immunol 10, 221, doi:10.3389/fimmu.2019.00221 (2019).

24 Burel, J. G. et al. An Integrated Workflow To Assess Technical and Biological Variability of Cell Population Frequencies in Human Peripheral Blood by Flow Cytometry. J Immunol 198, 1748–1758, doi:10.4049/jimmunol.1601750 (2017).

25 Arlehamn, C. L. et al. Transcriptional Profile of Tuberculosis Antigen-Specific T Cells Reveals Novel Multifunctional Features. J Immunol 193, 2931–2940, doi:10.4049/jimmunol.1401151 (2014).

26 Pomaznoy, M. et al. Quantitative and Qualitative Perturbations of CD8(+) MAITs in Healthy Mycobacterium tuberculosis-Infected Individuals. Immunohorizons 4, 292–307, doi:10.4049/immunohorizons.2000031 (2020).

27 Lindestam Arlehamn, C. S. et al. Immunological consequences of intragenus conservation of Mycobacterium tuberculosis T-cell epitopes. Proc Natl Acad Sci U S A 112, E147–155, doi:10.1073/pnas.1416537112 (2015).

28 Kleinnijenhuis, J. et al. Bacille Calmette-Guerin induces NOD2-dependent nonspecific protection from reinfection via epigenetic reprogramming of monocytes. Proc Natl Acad Sci U S A 109, 17537–17542, doi:10.1073/pnas.1202870109 (2012).

29 Verma, D. et al. Anti-mycobacterial activity correlates with altered DNA methylation pattern in immune cells from BCG-vaccinated subjects. Scientific reports 7, 12305, doi:10.1038/s41598-017-12110-2 (2017).

30 Houseman, E. A. et al. Reference-free deconvolution of DNA methylation data and mediation by cell composition effects. BMC Bioinformatics 17, 259, doi:10.1186/s12859-016-1140-4 (2016).

31 Chronister, W. D. et al. TCRMatch: Predicting T-Cell Receptor Specificity Based on Sequence Similarity to Previously Characterized Receptors. Front Immunol 12, 640725, doi:10.3389/fimmu.2021.640725 (2021).

32 Fletcher, H. A. Systems approaches to correlates of protection and progression to TB disease. Semin Immunol 39, 81–87, doi:10.1016/j.smim.2018.10.001 (2018).

33 Satti, I. & McShane, H. Current approaches toward identifying a correlate of immune protection from tuberculosis. Expert Rev Vaccines 18, 43–59, doi:10.1080/14760584.2019.1552140 (2019).

34 Barker, L., Hessel, L. & Walker, B. Rational approach to selection and clinical development of TB vaccine candidates. Tuberculosis (Edinb) 92 Suppl 1, S25–29, doi:10.1016/S1472-9792(12)70009-4 (2012).

35 Aranday Cortes, E., Kaveh, D., Nunez-Garcia, J., Hogarth, P. J. & Vordermeier, H. M. Mycobacterium bovis-BCG vaccination induces specific pulmonary transcriptome biosignatures in mice. PLoS One 5, e11319, doi:10.1371/journal.pone.0011319 (2010).

36 Hoft, D. F. et al. PO and ID BCG vaccination in humans induce distinct mucosal and systemic immune responses and CD4(+) T cell transcriptomal molecular signatures. Mucosal Immunol 11, 486–495, doi:10.1038/mi.2017.67 (2018).

37 Morgan, J. et al. Classical CD4 T cells as the cornerstone of antimycobacterial immunity. Immunol Rev 301, 10–29, doi:10.1111/imr.12963 (2021).

38 Caccamo, N. et al. Multifunctional CD4(+) T cells correlate with active Mycobacterium tuberculosis infection. Eur J Immunol 40, 2211–2220, doi:10.1002/eji.201040455 (2010).

39 Qiu, Z. et al. Multifunctional CD4 T cell responses in patients with active tuberculosis. Scientific reports 2, 216, doi:10.1038/srep00216 (2012).

40 Smith, S. G., Zelmer, A., Blitz, R., Fletcher, H. A. & Dockrell, H. M. Polyfunctional CD4 T-cells correlate with in vitro mycobacterial growth inhibition following Mycobacterium bovis BCG-vaccination of infants. Vaccine 34, 5298–5305, doi:10.1016/j.vaccine.2016.09.002 (2016).

41 Murray, R. A. et al. Bacillus Calmette Guerin vaccination of human newborns induces a specific, functional CD8+ T cell response. J Immunol 177, 5647–5651, doi:10.4049/jimmunol.177.8.5647 (2006).

42 Lindestam Arlehamn, C. S. et al. Memory T cells in latent Mycobacterium tuberculosis infection are directed against three antigenic islands and largely contained in a CXCR3+CCR6+ Th1 subset. PLoS Pathog 9, e1003130, doi:10.1371/journal.ppat.1003130 (2013).

43 Gela, A. et al. Effects of BCG vaccination on donor unrestricted T cells in humans. bioRxiv, 2021.2004.2029.441927, doi:10.1101/2021.04.29.441927 (2021).

44 Nathan, A. et al. Multimodally profiling memory T cells from a tuberculosis cohort identifies cell state associations with demographics, environment and disease. Nat Immunol 22, 781–793, doi:10.1038/s41590-021-00933-1 (2021).

45 Dijkman, K. et al. Prevention of tuberculosis infection and disease by local BCG in repeatedly exposed rhesus macaques. Nat Med 25, 255–262, doi:10.1038/s41591-018-0319-9 (2019).

46 Cadena, A. M. et al. Concurrent infection with Mycobacterium tuberculosis confers robust protection against secondary infection in macaques. PLoS Pathog 14, e1007305, doi:10.1371/journal.ppat.1007305 (2018).

47 Mahnke, Y. D., Brodie, T. M., Sallusto, F., Roederer, M. & Lugli, E. The who’s who of T-cell differentiation: human memory T-cell subsets. Eur J Immunol 43, 2797–2809, doi:10.1002/eji.201343751 (2013).

48 Sutherland, J. S., Adetifa, I. M., Hill, P. C., Adegbola, R. A. & Ota, M. O. C. Pattern and diversity of cytokine production differentiates between Mycobacterium tuberculosis infection and disease. Eur J Immunol 39, 723–729, doi:10.1002/eji.200838693 (2009).

49 Somoskovi, A. & Salfinger, M. Nontuberculous mycobacteria in respiratory infections: advances in diagnosis and identification. Clinics in laboratory medicine 34, 271–295, doi:10.1016/j.cll.2014.03.001 (2014).

50 Srivastava, S. & Ernst, J. D. Cutting edge: Direct recognition of infected cells by CD4 T cells is required for control of intracellular Mycobacterium tuberculosis in vivo. J Immunol 191, 1016–1020, doi:10.4049/jimmunol.1301236 (2013).

51 Anderson, K. G. et al. Intravascular staining for discrimination of vascular and tissue leukocytes. Nature protocols 9, 209–222, doi:10.1038/nprot.2014.005 (2014).

52 Sakai, S. et al. Cutting edge: control of Mycobacterium tuberculosis infection by a subset of lung parenchyma-homing CD4 T cells. J Immunol 192, 2965–2969, doi:10.4049/jimmunol.1400019 (2014).

53 Glanville, J. et al. Identifying specificity groups in the T cell receptor repertoire. Nature 547, 94–98, doi:10.1038/nature22976 (2017).

54 Song, I. et al. Broad TCR repertoire and diverse structural solutions for recognition of an immunodominant CD8(+) T cell epitope. Nat Struct Mol Biol 24, 395–406, doi:10.1038/nsmb.3383 (2017).

55 Sulzer, D. et al. T cells from patients with Parkinson’s disease recognize alpha-synuclein peptides. Nature 546, 656–661, doi:10.1038/nature22815 (2017).

56 Lindestam Arlehamn, C. S. et al. A Quantitative Analysis of Complexity of Human Pathogen-Specific CD4 T Cell Responses in Healthy M. tuberculosis Infected South Africans. PLoS Pathog 12, e1005760, doi:10.1371/journal.ppat.1005760 (2016).

57 da Silva Antunes, R. et al. Definition of Human Epitopes Recognized in Tetanus Toxoid and Development of an Assay Strategy to Detect Ex Vivo Tetanus CD4+ T Cell Responses. PLoS One 12, e0169086, doi:10.1371/journal.pone.0169086 (2017).

58 Corbett, A. J. et al. T-cell activation by transitory neo-antigens derived from distinct microbial pathways. Nature 509, 361–365, doi:10.1038/nature13160 (2014).

59 Trapnell, C., Pachter, L. & Salzberg, S. L. TopHat: discovering splice junctions with RNA-Seq. Bioinformatics 25, 1105–1111, doi:10.1093/bioinformatics/btp120 (2009).

60 Schmieder, R. & Edwards, R. Quality control and preprocessing of metagenomic datasets. Bioinformatics 27, 863–864, doi:10.1093/bioinformatics/btr026 (2011).

61 Li, H. et al. The Sequence Alignment/Map format and SAMtools. Bioinformatics 25, 2078–2079, doi:10.1093/bioinformatics/btp352 (2009).

62 Anders, S., Pyl, P. T. & Huber, W. HTSeq--a Python framework to work with high-throughput sequencing data. Bioinformatics 31, 166–169, doi:10.1093/bioinformatics/btu638 (2015).

63 Love, M. I., Huber, W. & Anders, S. Moderated estimation of fold change and dispersion for RNA-seq data with DESeq2. Genome Biol 15, 550, doi:10.1186/s13059-014-0550-8 (2014).

64 Benjamini, Y. & Hochberg, Y. Controlling the False Discovery Rate: A Practical and Powerful Approach to Multiple Testing. Journal of the Royal Statistical Society. Series B (Methodological) 57, 289–300 (1995).

65 Chen, J., Bardes, E. E., Aronow, B. J. & Jegga, A. G. ToppGene Suite for gene list enrichment analysis and candidate gene prioritization. Nucleic Acids Res 37, W305–311, doi:10.1093/nar/gkp427 (2009).

66 Schmiedel, B. J. et al. Impact of Genetic Polymorphisms on Human Immune Cell Gene Expression. Cell 175, 1701–1715 e1716, doi:10.1016/j.cell.2018.10.022 (2018).

67 Jones, E., Oliphant, T. & Peterson, P. SciPy: Open source scientific tools for Python. (2001).

68 Van Der Walt, S., Colbert, S. C. & Varoquaux, G. The NumPy array: a structure for efficient numerical computation. Computing in Science & Engineering 13, 22 (2011).

69 Seabold, S. & Perktold, J. in Proceedings of the 9th Python in Science Conference. 61 (Scipy).

70 Morris, T. J. et al. ChAMP: 450k Chip Analysis Methylation Pipeline. Bioinformatics 30, 428–430, doi:10.1093/bioinformatics/btt684 (2014).

71 Wilkinson, L. Exact and Approximate Area-Proportional Circular Venn and Euler Diagrams. IEEE Transactions on Visualization and Computer Graphics 18, 321–331, doi:10.1109/TVCG.2011.56 (2012).

72 Yu, G., Wang, L. G., Han, Y. & He, Q. Y. clusterProfiler: an R package for comparing biological themes among gene clusters. OMICS 16, 284–287, doi:10.1089/omi.2011.0118 (2012).

