## Supplemental figures and tables for "CD4+CCR6+ T cells dominate the BCG-induced transcriptional signature"

Cecilia S. Lindestam Arlehamn

**Supplementary material**

**Supplementary Figures**

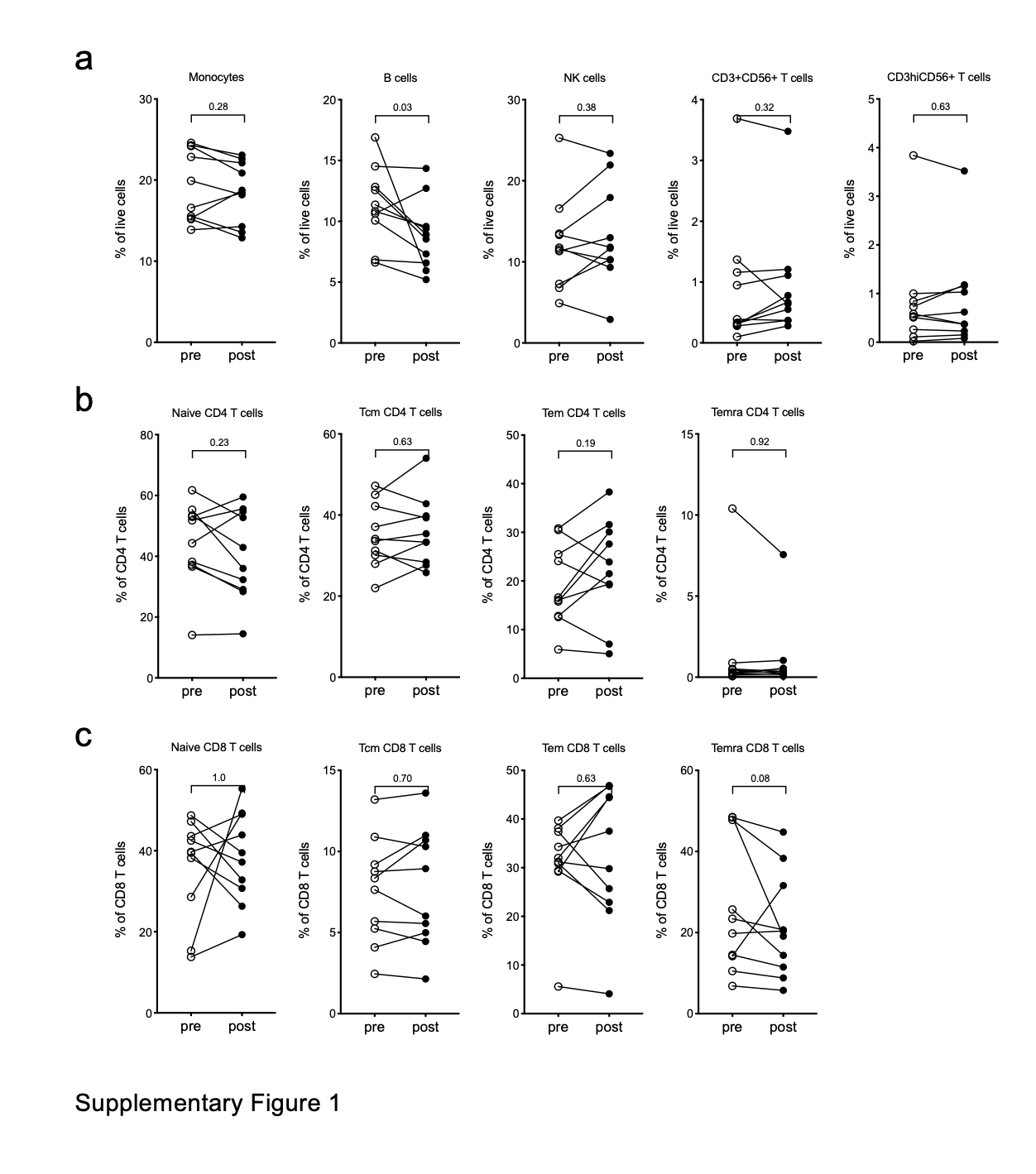

**SUPPLEMENTARY FIGURE 1. Cell subset frequencies pre- and post- BCG vaccination**. **a-c** Frequencies of cell subsets pre- (open circles) and post- (closed circles) BCG vaccination. Each point represents one participant, Wilcoxon matched pair signed rank test. **a** Major lymphocyte subsets not shown in figure 3. **b, c** Memory T cell subsets defined by CCR7 and CD45RA; Naïve cells (CD45RA+CCR7+), Tcm (CD45RA-CCR7+), Tem (CD45RA-CCR7-), and Temra (CD45RA+CCR7-). **b** CD4 T cell subsets. **c** CD8 T cell subsets.

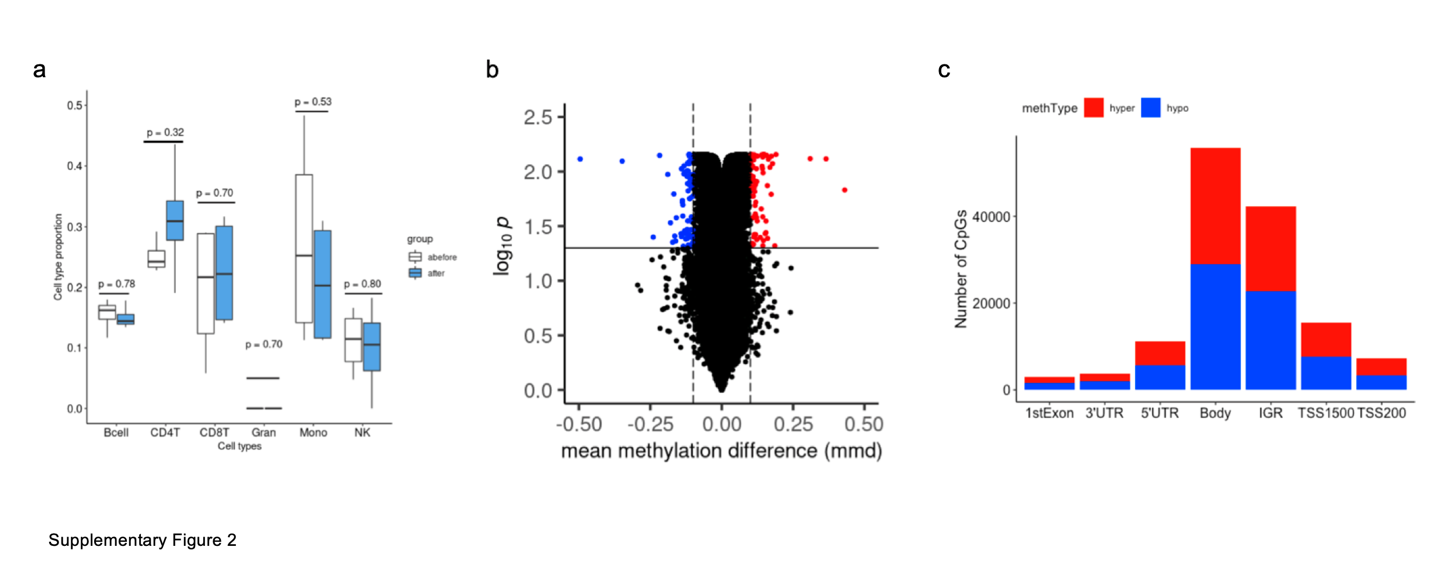

**SUPPLEMENTARY FIGURE 2. Characterization of the DNAm alterations**. **a**. Cell type frequency analysis applying the Houseman algorithm to the DNA methylation dataset of samples obtained pre- and post- BCG vaccination. The y-axis shows the frequency of cell types in the group of samples and the x-axis denotes the calculated cell type in the PBMC sample. Student’s t-test. **b**. Volcano plot illustrating the hyper- and hypomethylated CpGs. The x-axis represents the mean methylation difference (mmd). The y-axis denotes the significance level (p-value) at the logarithm scale. The solid horizontal line sets the threshold of p-value < 0.05 and the dashed vertical lines used for the cut-off value, mmd ≥ |0.1|. **c**. Number of CpGs present in different chromosomal locations for hypermethylated (red) and hypomethylated (blue) CpGs.

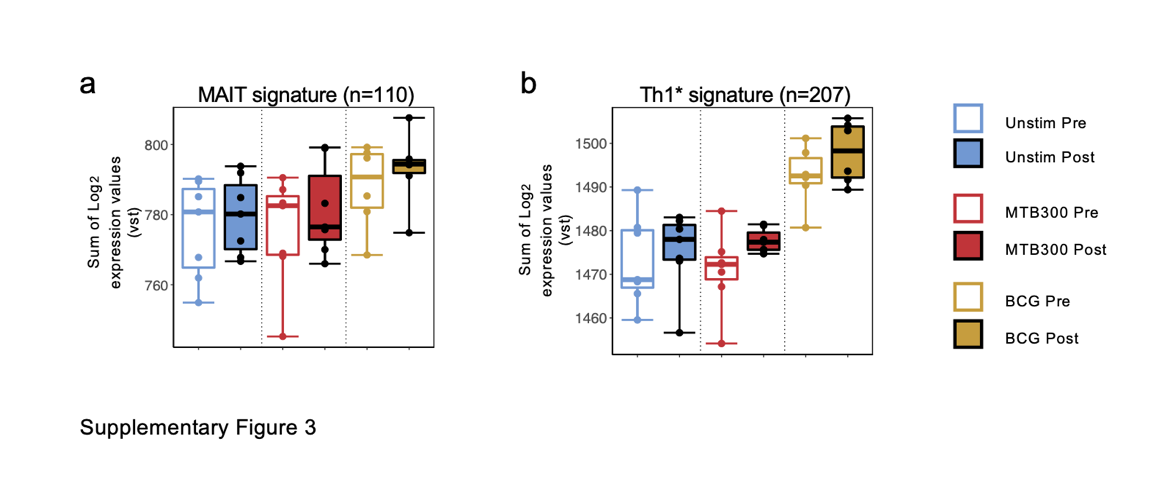

**SUPPLEMENTARY FIGURE 3**. **Cell subset-specific gene signatures pre- and post- BCG vaccination a, b** Boxplots of RNA-sequencing data depicting the sum of log_2_ expression values (variance stabilizing transformation, VST) for all genes in the **(a)** MAIT cell signature, and **(b)** Th1* cell signature. Empty and filled boxplot represent pre- and post-vaccination, respectively, and color represents stimulation condition. Each dot represents an individual donor.

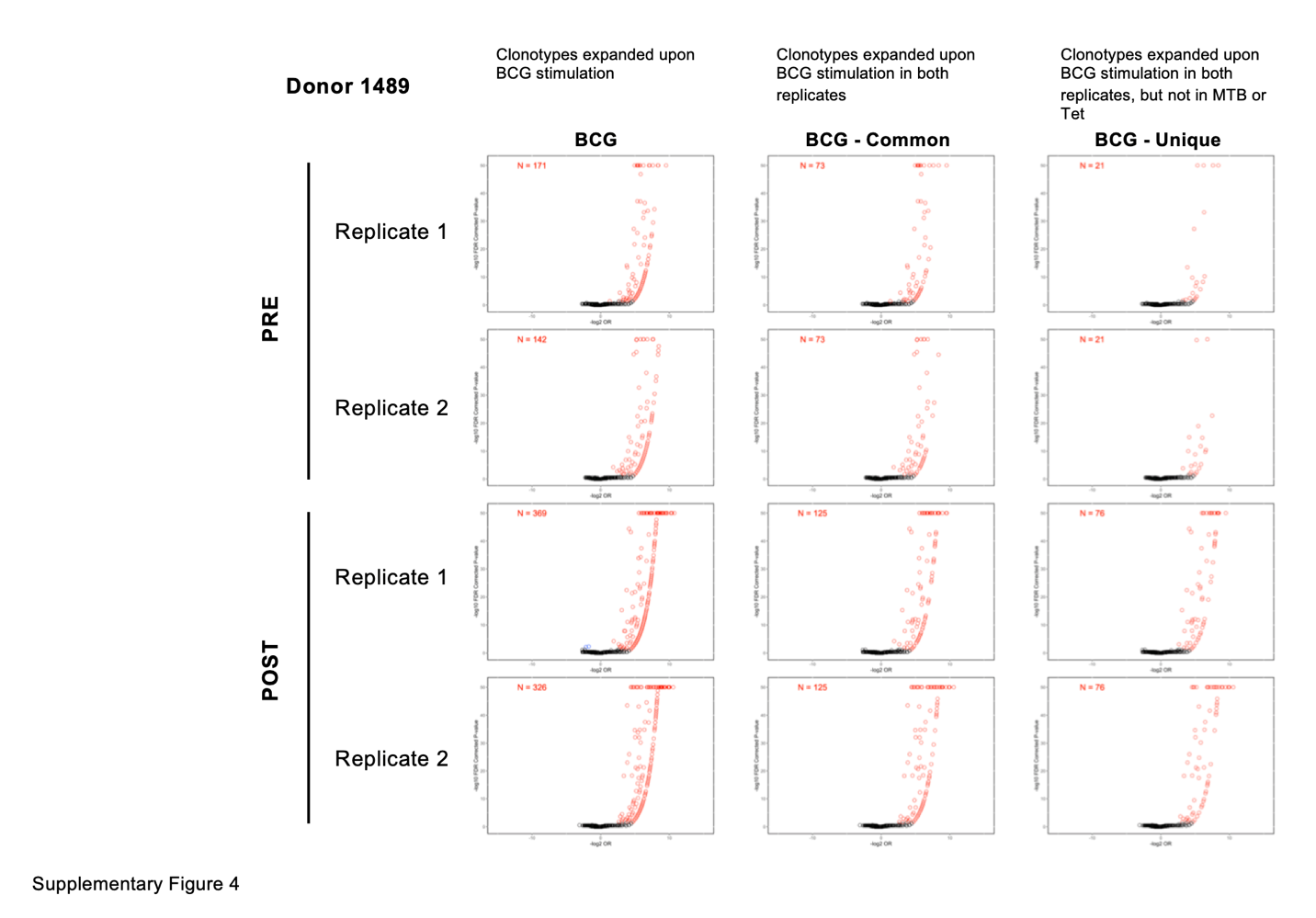

**SUPPLEMENTARY FIGURE 4.** Representative volcano plots showing clonotype expansion in response to BCG stimulation. Analysis from Donor 1489 is shown. Column 1 indicates clonotypes expanded upon BCG stimulation in each replicate, pre- and post- vaccination. Column 2 indicates only those clonotypes that expanded upon BCG stimulation in both replicates, pre- and post- vaccination. Column 3 further indicates a subset to show only those clonotypes that expanded upon BCG stimulation in both replicates, but not in MTB300 or Tetanus stimulation, pre- and post- vaccination.

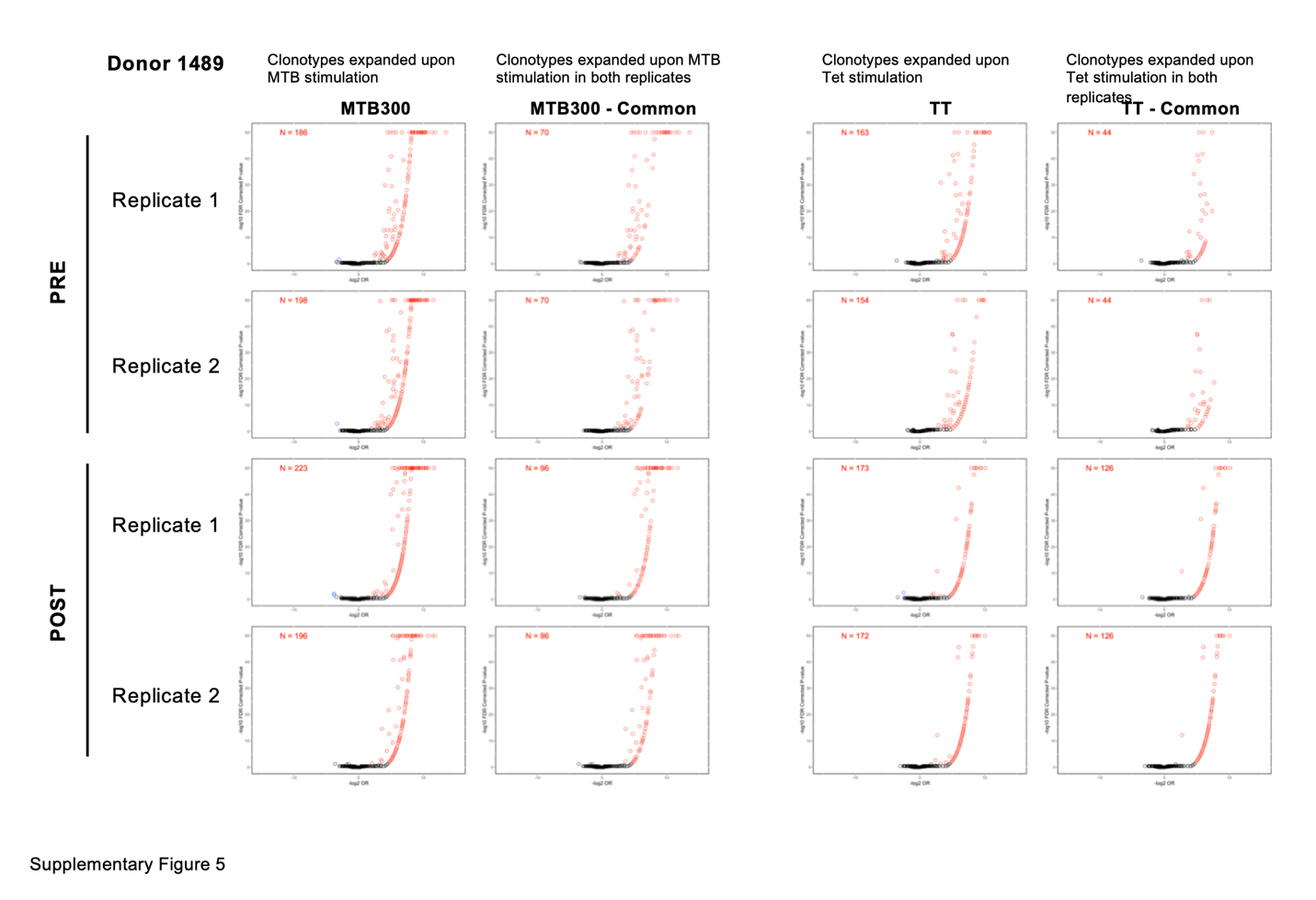

**SUPPLEMENTARY FIGURE 5**. Representative volcano plots showing clonotype expansion in response to MTB300 and Tetanus stimulation. Analysis from Donor 1489 is shown. Column 1 indicates clonotypes expanded upon MTB300 stimulation in each replicate, pre- and post- vaccination. Column 2 indicates only those clonotypes that expanded upon MTB300 stimulation in both replicates, pre- and post- vaccination. Column 3 indicates clonotypes expanded upon Tetanus stimulation in each replicate, pre- and post- vaccination. Column 4 indicates only those clonotypes that expanded upon Tetanus stimulation in both replicates, pre- and post- vaccination.

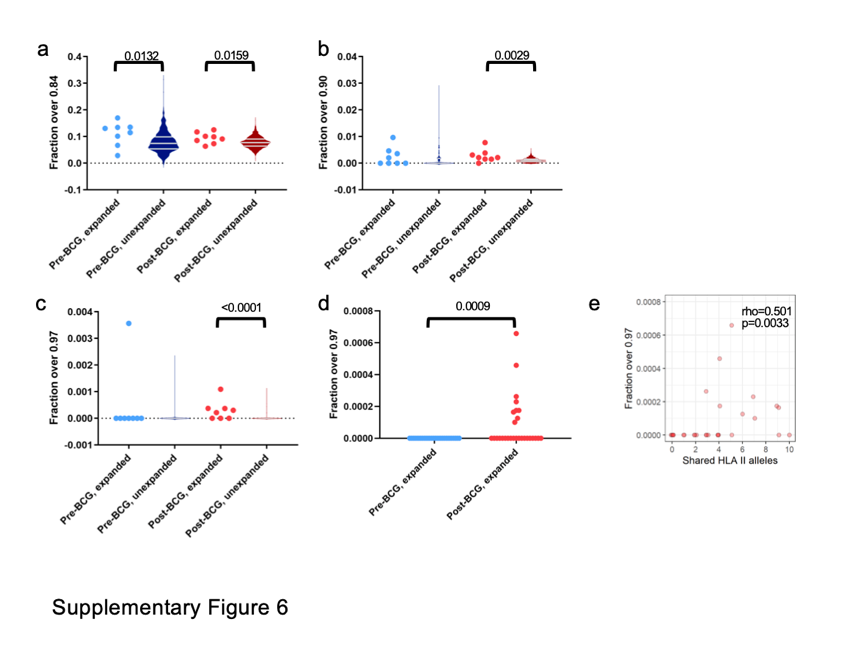

**SUPPLEMENTARY FIGURE 6. Comparison of TCRMatch scores within and across T cell repertoires from individuals pre- and post-BCG vaccination. a-c** Comparison of CDR3β similarity within individuals before and after BCG vaccination and expanded or unexpanded in response to BCG stimulus in vitro, as assessed by the fraction of scores (**a**) >0.84, (**b**) >0.90, (**c**) >0.97. 100 random samplings of each individual’s unexpanded TCRs are represented by violin plots. Gray lines indicate first, second and third quartiles. One-tailed Mann-Whitney test. **d** Comparison of CDR3β similarity in expanded T cells across individuals pre- and post-BCG vaccination. One-tailed Mann-Whitney test. **e** Comparison of the fraction of TCRMatch scores >0.97 between two individuals and the number of HLA class II alleles shared by the individuals. One-tailed Spearman test.

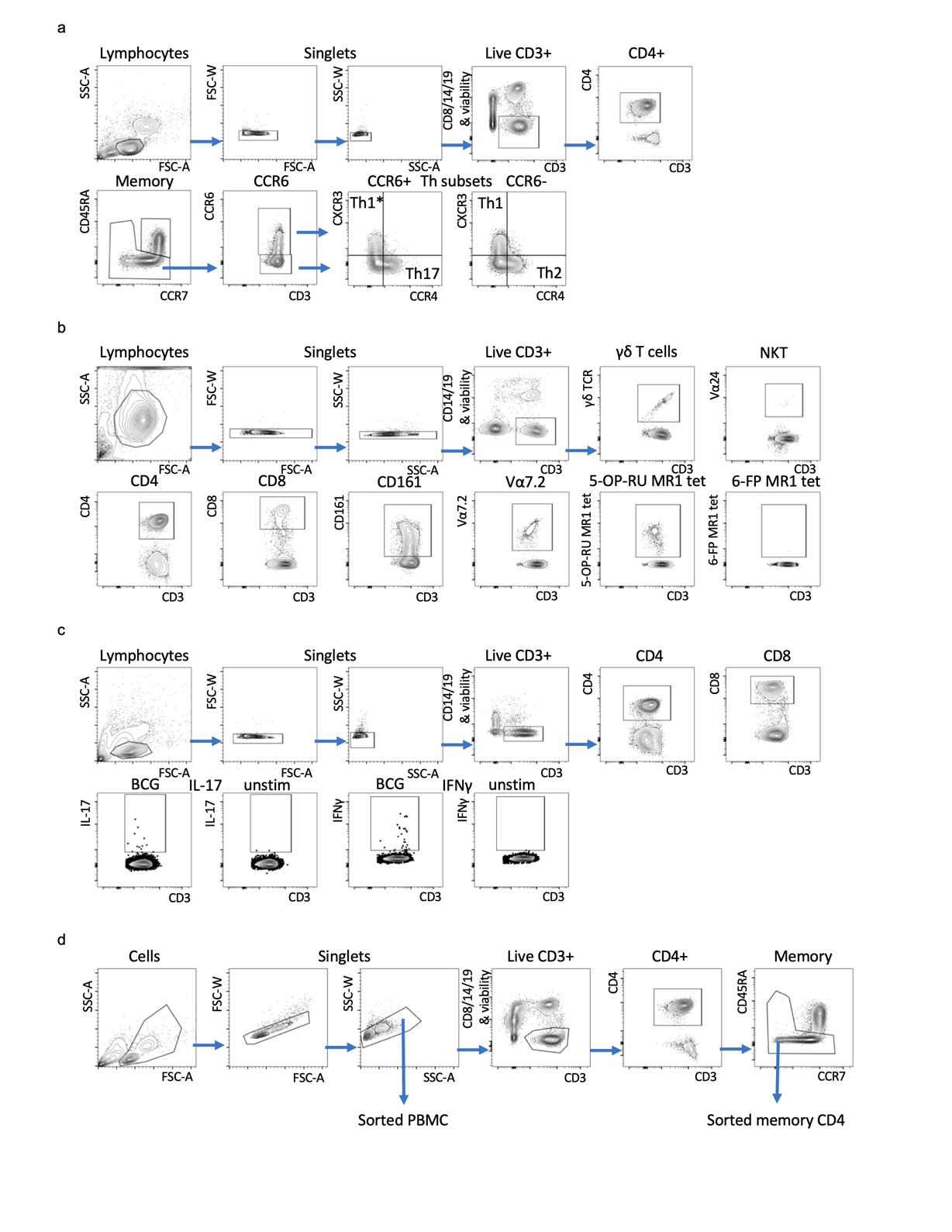

**SUPPLEMENTARY FIGURE 7. Gating strategies for flow cytometry analyses and cell sorting. a** Th-subset gating, **b** non-conventional T cells, MAITs were defined as MR1 5-OP-RU tetramer+CD4-Vα24-γδTCR-Vα7.2+CD161+. A representative control 6-FP MR1 tetramer stain is shown for comparison. **c** Intracellular cytokine staining. BCG stimulated and unstimulated samples for both IL-17 and IFNγ is shown. **d** Cell sorting of PBMC (excluding doublets) and live memory CD4 T cells.

**Supplementary Tables**

Supplementary table 1-4 provided as excel files

**SUPPLEMENTARY TABLE 1** Differentially expressed genes identified in the different comparisons for PBMC and CD4 memory T cells.

**SUPPLEMENTARY TABLE 2** Pathway enrichment for upregulated genes in PBMCs and CD4 memory T cells.

**SUPPLEMENTARY TABLE 3** MAIT and Th1* signatures

**SUPPLEMENTARY TABLE 4** TCR repertoire dataset

**SUPPLEMENTARY TABLE 5** Number of clonotypes per subject

|  | **Subject** | **Total Clonotypes** | | **Clonotypes expanded upon BCG stimulation** | | **Clonotypes expanded in both replicates** | | | | |
| --- | --- | --- | --- | --- | --- | --- | --- | --- | --- | --- |
|  |  | **Replicate 1** | **Replicate 2** | **Replicate 1** | **Replicate 2** | **BCG stim** | **BCG stim - tet** | **BCG stim only** | **MTB300 stim only** | **BCG and MTB300 stim** |
| **PRE** | 1489 | 30544 | 24621 | 171 | 142 | 73 | 36 | 30 | 33 | 6 |
|  | 1491 | 18875 | 17343 | 183 | 157 | 54 | 23 | 18 | 19 | 5 |
|  | 1492 | 4616 | 5515 | 105 | 209 | 22 | 20 | 19 | 13 | 1 |
|  | 1497 | 5597 | 6524 | 174 | 184 | 68 | 44 | 32 | 35 | 12 |
|  | 1499 | 9248 | 8944 | 205 | 147 | 84 | 62 | 46 | 27 | 16 |
|  | 1500 | 10476 | 12232 | 369 | 382 | 90 | 55 | 40 | 30 | 15 |
|  | 1502 | 13174 | 17269 | 235 | 289 | 55 | 40 | 34 | 34 | 6 |
|  | 2466 | 7930 | 11635 | 89 | 85 | 41 | 25 | 15 | 27 | 10 |
| **POST** | 1489 | 22744 | 19449 | 369 | 326 | 125 | 98 | 82 | 49 | 16 |
|  | 1491 | 17607 | 14725 | 363 | 298 | 131 | 79 | 59 | 48 | 20 |
|  | 1492 | 9533 | 12276 | 357 | 563 | 109 | 92 | 74 | 62 | 18 |
|  | 1497 | 6756 | 6840 | 299 | 277 | 117 | 85 | 68 | 85 | 17 |
|  | 1499 | 5408 | 5296 | 288 | 301 | 147 | 116 | 97 | 47 | 19 |
|  | 1500 | 12347 | 10739 | 602 | 658 | 170 | 145 | 127 | 66 | 18 |
|  | 1502 | 9916 | 11308 | 590 | 520 | 155 | 126 | 103 | 79 | 23 |
|  | 2466 | 7042 | 4341 | 225 | 232 | 98 | 76 | 61 | 41 | 15 |

**SUPPLEMENTARY TABLE 6** GLIPH analysis within and across donors to identify similarities in clonotypes that expanded pre- and post- vaccination, upon BCG stimulation in both replicates, but not in MTB300 or Tetanus stimulation.

|  | **Subject** | **Clonotypes expanded in** | | | **GLIPH analysis** | | | | |
| --- | --- | --- | --- | --- | --- | --- | --- | --- | --- |
|  |  | **Pre** | **Post** | **Common in**  **Pre & Post** | **3-mers** | **4-mers** | **Convergence groups** | **Global**  **convergences** | **Local**  **convergences** |
| Similarity between Pre and Post within each donor in BCG stim only | 2466 | 9 | 55 | 6 | - | - | 3 groups with 2 peptides each | 3 | - |
|  | 1502 | 20 | 89 | 14 | - | - | 7 groups with 2 peptides each | 7 | - |
|  | 1500 | 27 | 114 | 13 | - | - | 2 groups with 2 peptides each | 2 | - |
|  | 1499 | 12 | 63 | 34 | 2 | - | 1 group with 6 peptides  2 groups with 5 peptides each  3 groups with 2 peptides each | 23 | 20 |
|  | 1497 | 19 | 55 | 13 | 1 | - | 1 group with 5 peptides  4 groups with 2 peptides each | 7 | 6 |
|  | 1492 | 14 | 69 | 5 | - | - | 1 group with 5 peptides  3 groups with 2 peptides each | 4 | 6 |
|  | 1491 | 11 | 52 | 7 | - | - | - | - | - |
|  | 1489 | 13 | 65 | 17 | - | - | 1 group with 2 peptides | 1 | - |

**SUPPLEMENTARY TABLE 7** HLA alleles expressed by each subject

| **Subject** | **A** | **B** | **C** | **DPB1** | **DQA1** | **DQB1** | **DRB1** | **DRB3/4/5** |
| --- | --- | --- | --- | --- | --- | --- | --- | --- |
| 1489 | A*01:01 / A*02:01 | B*08:01 / B*15:01 | C*01:02 / C*07:01 | DPB1*02:01 / DPB1*04:01 | DQA1*04:01 / DQA1*05:01 | DQB1*02:01 / DQB1*04:02 | DRB1*03:01 / DRB1*08:01 | DRB3*01:01 / |
| 1490 | A*02:01 / A*03:01 | B*07:02 / B*44:05 | C*02:02 / C*07:02 | DPB1*03:01 / DPB1*11:01 | DQA1*02:01 / DQA1*03:01 | DQB1*02:02 / DQB1*03:02 | DRB1*04:04 / DRB1*07:01 | DRB4*01:01 / DRB4*01:01 |
| 1491 | A*02:01 / A*68:01 | B*15:01 / B*27:05 | C*02:02 / C*03:03 | DPB1*02:01 / DPB1*04:01 | DQA1*03:01 / DQA1*03:01 | DQB1*03:01 / DQB1*03:02 | DRB1*04:01 / DRB1*04:01 | DRB4*01:01 / DRB4*01:01 |
| 1492 | A*02:01 / A*32:01 | B*15:01 / B*44:02 | C*04:01 / C*05:01 | DPB1*03:01 / DPB1*20:01 | DQA1*03:01 / DQA1*04:01 | DQB1*03:02 / DQB1*04:02 | DRB1*04:01 / DRB1*08:01 | DRB4*01:01 / |
| 1493 | A*03:01 / A*03:01 | B*35:01 / B*56:01 | C*01:02 / C*04:01 | DPB1*04:01 / DPB1*04:02 | DQA1*01:01 / DQA1*01:02 | DQB1*05:01 / DQB1*06:02 | DRB1*01:01 / DRB1*15:01 | DRB5*01:01 / |
| 1494 | A*01:01 / A*02:01 | B*15:01 / B*55:01 | C*03:03 / C*03:04 | DPB1*04:01 / DPB1*04:02 | DQA1*03:01 / DQA1*05:01 | DQB1*03:01 / DQB1*03:02 | DRB1*04:01 / DRB1*11:04 | DRB3*02:02 / DRB4*01:01 |
| 1495 | A*25:01 / A*26:01 | B*18:01 / B*51:07 | C*12:03 / C*14:02 | DPB1*04:01 / DPB1*19:01 | DQA1*01:02 / DQA1*01:03 | DQB1*06:03 / DQB1*06:09 | DRB1*13:01 / DRB1*13:02 | DRB3*02:02 / DRB3*03:01 |
| 1496 | A*02:01 / A*33:03 | B*40:23 / B*50:01 | C*03:04 / C*06:02 | DPB1*04:02 / DPB1*14:01 | DQA1*03:01 / DQA1*03:01 | DQB1*03:01 / DQB1*03:02 | DRB1*04:01 / DRB1*09:01 | DRB4*01:01 / DRB4*01:01 |
| 1497 | A*01:01 / A*02:01 | B*44:02 / B*44:02 | C*05:01 / C*05:01 | DPB1*04:01 / DPB1*04:01 | DQA1*01:02 / DQA1*05:01 | DQB1*03:01 / DQB1*06:02 | DRB1*12:01 / DRB1*15:01 | DRB3*02:02 / DRB5*01:01 |
| 1498 | A*03:01 / A*68:01 | B*44:02 / B*57:01 | C*06:02 / C*07:12 | DPB1*01:01 / DPB1*04:01 | DQA1*01:01 / DQA1*02:01 | DQB1*03:03 / DQB1*05:01 | DRB1*01:01 / DRB1*07:01 | DRB4*01:01 / |
| 1499 | A*01:01 / A*02:01 | B*08:01 / B*15:01 | C*03:04 / C*07:01 | DPB1*04:01 / DPB1*04:01 | DQA1*03:01 / DQA1*05:01 | DQB1*02:01 / DQB1*03:02 | DRB1*03:01 / DRB1*04:01 | DRB3*01:01 / DRB4*01:01 |
| 1500 | A*03:01 / A*03:01 | B*07:02 / B*07:02 | C*07:02 / C*07:02 | DPB1*03:01 / DPB1*03:01 | DQA1*01:01 / DQA1*01:02 | DQB1*05:01 / DQB1*06:02 | DRB1*10:01 / DRB1*15:01 | DRB5*01:01 / |
| 1501 | A*02:01 / A*02:01 | B*15:01 / B*40:01 | C*03:03 / C*03:04 | DPB1*02:01 / DPB1*03:01 | DQA1*03:01 / DQA1*03:01 | DQB1*03:01 / DQB1*03:02 | DRB1*04:01 / DRB1*04:07 | DRB4*01:01 / DRB4*01:01 |
| 1502 | A*02:01 / A*03:01 | B*08:01 / B*44:02 | C*05:01 / C*07:01 | DPB1*04:01 / DPB1*04:01 | DQA1*01:02 / DQA1*03:01 | DQB1*03:01 / DQB1*06:02 | DRB1*04:01 / DRB1*15:01 | DRB4*01:01 / DRB5*01:01 |
| 1503 | A*02:01 / A*03:01 | B*07:02 / B*15:01 | C*03:03 / C*07:02 | DPB1*02:01 / DPB1*04:01 | DQA1*01:02 / DQA1*03:01 | DQB1*03:02 / DQB1*06:04 | DRB1*04:01 / DRB1*13:02 | DRB3*03:01 / DRB4*01:01 |
| 1504 | A*01:01 / A*02:01 | B*08:01 / B*15:01 | C*03:04 / C*07:01 | DPB1*03:01 / DPB1*04:01 | DQA1*03:01 / DQA1*05:01 | DQB1*02:01 / DQB1*03:02 | DRB1*03:01 / DRB1*04:01 | DRB3*01:01 / DRB4*01:01 |
| 2466 | A*02:01 / A*02:01 | B*40:01 / B*44:02 | C*03:04 / C*05:01 | DPB1*02:01 / DPB1*04:02 | DQA1*02:01 / DQA1*03:01 | DQB1*02:02 / DQB1*03:01 | DRB1*04:01 / DRB1*07:01 | DRB4*01:01 / DRB4*01:01 |
